# Widespread associations between grey matter structure and the human phenome

**DOI:** 10.1101/696864

**Authors:** Baptiste Couvy-Duchesne, Lachlan T. Strike, Futao Zhang, Yan Holtz, Zhili Zheng, Kathryn E. Kemper, Loic Yengo, Olivier Colliot, Margaret J. Wright, Naomi R. Wray, Jian Yang, Peter M. Visscher

**Affiliations:** Institute for Molecular Bioscience, the University of Queensland, 4072 St Lucia, QLD, Australia; Queensland Brain Institute, the University of Queensland, 4072 St Lucia, QLD, Australia; Institute for Advanced Research, Wenzhou Medical University, Wenzhou, Zhejiang 325027, China; Inria, ARAMIS Project-team, 75013, Paris, France; Institut du Cerveau et de la Moelle épinière, 75013, Paris, France; Inserm, U1127, 75013, Paris, France; CNRS, UMR 7225, 75013, Paris, France; Sorbonne Universitee, 75013, Paris, France; Centre for Advanced Imaging, the University of Queensland, 4072 St Lucia, QLD, Australia

## Abstract

The recent availability of large-scale neuroimaging cohorts (here the UK Biobank [UKB] and the Human Connectome Project [HCP]) facilitates deeper characterisation of the relationship between phenotypic and brain architecture variation in humans. We tested the association between 654,386 vertex-wise measures of cortical and subcortical morphology (from T1w and T2w MRI images) and behavioural, cognitive, psychiatric and lifestyle data. We found a significant association of grey-matter structure with 58 out of 167 UKB phenotypes spanning substance use, blood assay results, education or income level, diet, depression, being a twin as well as cognition domains (UKB discovery sample: N=9,888). Twenty-three of the 58 associations replicated (UKB replication sample: N=4,561; HCP, N=1,110). In addition, differences in body size (height, weight, BMI, waist and hip circumference, body fat percentage) could account for a substantial proportion of the association, providing possible insight into previous MRI case-control studies for psychiatric disorders where case status is associated with body mass index. Using the same linear mixed model, we showed that most of the associated characteristics (e.g. age, sex, body size, diabetes, being a twin, maternal smoking, body size) could be significantly predicted using all the brain measurements in out-of-sample prediction. Finally, we demonstrated other applications of our approach including a Region Of Interest (ROI) analysis that retain the vertex-wise complexity and ranking of the information contained across MRI processing options.

**Highlights:**

- Our linear mixed model approach unifies association and prediction analyses for highly dimensional vertex-wise MRI data
- Grey-matter structure is associated with measures of substance use, blood assay results, education or income level, diet, depression, being a twin as well as cognition domains
- Body size (height, weight, BMI, waist and hip circumference) is an important source of covariation between the phenome and grey-matter structure
- Grey-matter scores quantify grey-matter based risk for the associated traits and allow to study phenotypes not collected
- The most general cortical processing (“fsaverage” mesh with no smoothing) maximises the brain-morphometricity for all UKB phenotypes

## 1. Introduction

The field of MRI studies is at a turning point owing to the recent availability of large data sets to researchers, including the UKB (Miller et al., 2016) and HCP (Van Essen et al., 2013; Van Essen et al., 2012b) samples. These datasets promote not only the replication of previous findings, but also expand the range of phenotypes available for study (e.g. psychiatric symptoms and lifestyle factors). In addition, such data sets can offer insights into the brain markers that may be shared between phenotypes, helping to draw new links between brain and behaviour. Finally, these community samples can complement the typical case-control paradigm by identifying confounders of MRI analyses or by studying related traits (e.g. cognition domains relevant in Alzheimer’s disease).

Here, we introduce a set of analyses that leverages large sample sizes to fully exploit the spatial resolution of MRI images using linear mixed models (LMM) implemented in the OSCA software tool (Zhang et al., 2019). Our high-resolution approach (i.e. vertex-wise morphological measures) has the advantage of retaining all the brain complexity data of current MRI acquisitions rather than relying on prior-based data reduction techniques (e.g. the region-of-interest [ROI] approach), and allows for the elucidation of precise brain-phenotype associations.

Specifically, we used an efficient implementation of LMMs to estimate the multivariate correlation of 600,000+ cortical and subcortical measurement at vertices extracted from T1 weighted (T1w) and T2 weighted (T2w) MRI images with a phenotype of interest (previously coined morphometricity (Sabuncu et al., 2016), here we prefer the more specific brain-morphometricity). We extended this framework to also estimate the proportion of variance in a trait associated with the vertex-wise data from specific brain features, hemispheres and regions of interest. We further introduce multi-trait LMMs that can further quantify shared brain-morphometricity (grey-matter correlation) between traits, reflecting causal, bi-directional or confounded relationships. In addition, we show how LMMs can estimate the joint effects of all brain features on a trait to construct a trait predictor from brain features (grey-matter score) that can be applied and tested in an independent sample. As such, our approach unifies association studies and prediction analyses, in order to unravel the brain-phenome relationships (Rosenberg et al., 2018).

We analysed two of the largest MRI datasets available (UKB [split into discovery N=9,888 and replication N=4,561] and HCP [N=1,110]) and considered a wide range of phenotypes spanning demographics, blood cell composition, diet, psychiatric and traumatic history, physical capacities or substance use. We have released our image processing and analysis software/scripts as well as all summary statistics to facilitate replication and re-use of the results.

## 2. Materials and Methods

### 2.1. UK Biobank (UKB) sample

#### 2.1.1. Participants recruitment, inclusion and exclusion criteria

The UKB participants were unselected volunteers from the United Kingdom (Sudlow et al., 2015). Participants who had participated in the baseline UKB data collection were invited to undergo the imaging study if they lived within travelling distance of the imaging centre. Exclusion criteria were limited to: presence of metal implant, recent surgery and health conditions problematic for MRI imaging (e.g. hearing, breathing problems or extreme claustrophobia) (Miller et al., 2016).

#### 2.1.2. T1 and T2 FLAIR image collection

MRI images were collected in Cheadle (greater Manchester) using a 3T Siemens Skyra machine (software platform VD13) and a 32-channel head coil (Miller et al., 2016). The T1 weighted (T1w) images were acquired over 4:54 minutes, voxel size 1.0×1.0×1.0mm, matrix of 208×256×256mm, using a 3D MPRAGE sequence (Mugler and Brookeman, 1990), sagittal orientation of slice acquisition, R=2 (in plane acceleration factor), TI/TR=880/2000 ms (Miller et al., 2016). The T2 FLAIR acquisition lasted 5:52 minutes, voxel size 1.05×1.0×1.0 mm, matrix of 192×256×256 voxels, 3D SPACE sequence (Mugler et al., 2000), sagittal orientation, R=2, partial Fourier 7/8, fat saturated, TI/TR=1800/5000ms, elliptical (Miller et al., 2016).

#### 2.1.3. Image processing

We processed the T1w and T2 FLAIR images using the ENIGMA (Thompson et al., 2014) protocols for cortical surface and thickness (Stein et al., 2012) as well as subcortical structure (Gutman et al., 2013; Gutman et al., 2012). When both T1w and T2 FLAIR were available for a participant, we processed them together to enhance the tissue segmentation in FreeSurfer 6.0 (Fischl, 2012), hence a more precise skull stripping and pial surfaces definition. When the T2 FLAIR was not acquired or not usable, we processed the T1w image by itself. We retained the full image information by using the (fsaverage) vertex-wise level data in the cortical surface and thickness analyses. This corresponded to 149,960 cortical vertices in the left hemisphere and 149,933 in the right hemisphere, for each modality. In addition, we extracted subcortical radial thickness and log Jacobian determinant (that measures surface deformation from a template, somewhat analogous to a relative surface area (Roshchupkin et al., 2016)) for 27,300 vertices per hemisphere mapping 7 subcortical volumes (hippocampus, putamen, amygdala, thalamus, caudate, pallidum and accumbens) (Gutman et al., 2013). Overall, the imaging data used in the analyses comprised 654,386 vertex measurements per individual: 299,893 describing cortical thickness, another 299,893 for cortical surface area, 27,300 for subcortical thickness and 27,300 for subcortical curvature.

For comparison with previous ENIGMA publications, we also extracted cortical thickness and surface area of 34 cortical regions delimited by the Desikan atlas (Desikan et al., 2006; Fischl et al., 2004), as described on the ENIGMA website. To further the comparison of processing options, we extracted cortical measurements from smoothed fsaverage meshes (fwhm 5, 10, 15, 20 and 25mm) as well as (unsmoothed) coarser meshes provided by FreeSurfer: fsaverage6 (149,091 vertices for both hemispheres and modalities), fsaverage5 (37,455 vertices), fsaverage4 (9,457 vertices) and fsaverage3 (2,414).

#### 2.1.4. Discovery Sample description

At the time of download (July 2017), T1w images were available for 10,102 participants of the UK Biobank (UKB) project. None of the participants had withdrawn consent after the data was collected. We excluded 175 participants with T1w images labelled as unusable by the UKB, leaving 9,928 MRI scans to process. T2 FLAIR images were available for 9,755 of those. The FreeSurfer processing failed or did not complete within 48 hours for a handful of participants: 37 for cortical processing, 19 for subcortical, including 17 for whom both processing failed. For simplicity, we chose not to re-run image processing on these participants as their exclusion should have a minimal impact on the results obtained from the full sample. Excluded individuals are described in **Dataset S1**. Our final sample comprised 9,890 participants with usable cortical data, 9,908 with subcortical data and 9,888 with both cortical and subcortical data. This sample consisted of 9,888 adults aged 62.5 on average (SD=7.5, range 44.6–79.6) and comprised 52.4% of female participants. We excluded 391 participants with extreme brains (outliers) or likely to have a large effect on the analyses (see **Appendix S1** for details of QC and **Dataset S1** for description of the excluded participants).

#### 2.1.5. Variables used

We included 168 variables grouped in several categories: demographics, cognition, physical test, psychiatry, recent feelings, stress and traumas, substance use, miscellaneous, brain measurements, blood assay and diet (see **Dataset S2** for details). When longitudinal observations were available for a participant, we used the one collected as part of the imaging assessment (when available) or the closest in time.

#### 2.1.6. Replication Sample description

Replication data set was downloaded in May 2018 and consisted in an additional 4,942 participants with a T1w image. Image processing and phenotype selection were identical to that of the discovery sample. This led to the exclusion of 381 participants whose processing failed and 238 excluded from QC. The final sample (N=4,323) included in the replication analysis was on average 63.1 years old (SD=7.46, range 46.1-80.3) with 52.1% of females. The age difference between discovery and replication sample was small but significant (p=9.02e-7). See **Dataset S1** for a full description of replication participants (final, QCed and failed processing) in addition to a comparison of the discovery and replication samples.

### 2.2. Human Connectome Project (HCP) sample

#### 2.2.1. Participants recruitment, inclusion and exclusion criteria

HCP participants were recruited from ongoing longitudinal studies of the Missouri Family Study (Edens et al., 2010; Sartor et al., 2011) and had to be between 22 and 35 years of age. Inclusion and exclusion criteria have been described previously (Van Essen et al., 2012b).

#### 2.2.2. T1 and T2 weighted image collection

T1w and T2 weighted (T2w) images were collected at the Washington University (St Louis, Missouri) on a 3T Siemens Skyra scanner using a standard 32-channel head coil (Van Essen et al., 2013; Van Essen et al., 2012b). Two T1w images were acquired, each over 7 minutes and 40 seconds with a voxel size of 0.7×0.7×0.7mm, matrix/FOV of 224×224×224mm using a 3D MPRAGE sequence (Mugler and Brookeman, 1990), TR/TE/TI=2400/2.14/1000ms, flip angle 8degrees, R=2, sagittal orientation of slice acquisition (Glasser et al., 2013). Similarly, two T2w images were acquired over 8:24 min each, voxel size 0.7×0.7×0.7mm, matrix of 224×224×224mm, 3DSPACE sequence (Mugler et al., 2000), sagittal orientation, R=2, TR/TE=3200/565, no fat suppression pulse.

#### 2.2.3. Image processing

The HCP team (Glasser et al., 2013; Marcus et al., 2013; Van Essen et al., 2012a) pre-processed the structural scans to facilitate scan comparison across individuals, removing spatial artefacts and improve T1w and T2w alignment using FSL (Jenkinson et al., 2002; Jenkinson et al., 2012) and FreeSurfer (Fischl, 2012). When both passed HCP quality control (QC), T1w and T2w images they processed them together in FreeSurfer 6.0 (Fischl, 2012), otherwise data extraction relied on a single scan (Glasser et al., 2013). Participants with poor quality T1w and T2w scans were re-imaged (Glasser et al., 2013). Cortical processing (recon-all procedure in FreeSurfer) was also performed by the HCP team and included down sampling to 1mm size voxels and 256×256×256 matrix, aided registration using customised brain mask, and two manual steps performed outside of the recon-all procedure to enhance white matter and pial reconstruction (Glasser et al., 2013). We downloaded the processed images (Marcus et al., 2011) and performed ENIGMA shape analysis (Gutman et al., 2013; Gutman et al., 2012) to extract vertex-wise measurements of the subcortical thickness and curvature. As for the UKB sample, a total of 654,386 vertex measurements were extracted for each individual. We excluded 24 outliers with extreme brains or likely to bias the analyses (see **Appendix S1** and **Dataset S2** for description of excluded participants).

#### 2.2.4. Sample description

As per the HCP “1200 Subjects data release” (1^st^ of March 2017), 1,113 participants were scanned on the 3T MRI and underwent extensive behavioural testing. Participants were mostly (54.4%) females and were 28.8 years old on average (SD=3.7, range 22–37). The sample comprised 455 twins (41.0%), 286 monozygotic twins (138 complete pairs) and 169 dizygotic twins (78 complete pairs). In addition, siblings and half siblings of twins were also recruited which resulted in only 445 distinct families in the sample.

#### 2.2.5. Variables used

For the HCP sample, we included 161 variables, some of which were also available in the UKB (e.g. demographics, cognition, physical assessment, blood assay or psychiatry). We also included interesting variables only present in the HCP sample: personality, emotion, in depth mental health assessment (Semi-Structured Assessment for the Genetics of Alcoholism (SSAGA) and Adult Self Report (ASR) (Achenbach, 2009; Achenbach et al., 2003)), detailed cognition, Pittsburgh sleep index (PSQI) (Buysse et al., 1989), or results from the urine drug tests (see **Dataset S2**).

### 2.3. Variance component analyses and brain relatedness matrix calculation

#### 2.3.1. The linear mixed model

We aimed to estimate the variance of a trait accounted for by brain features, which Sabuncu et al., called “morphometricity” (Sabuncu et al., 2016). To do so we consider the following linear mixed model that allows estimating the association between a phenotype and M vertices even when M is greater than the sample size (N):

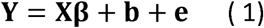

where **Y**_N,1_ is the phenotype considered with N the number of observations, **X**_N,c_ is a matrix of c covariates (as such does not include any vertex variable), **β**_c,1_ is a vector of fixed effects, **b** is a random effect with 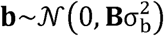 and **e** is the error term with 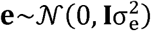. In this formulation **1**_N,N_ is the identity matrix as we assume the error terms to be independent and identically distributed. **B**_N,N_ is a matrix of variance-covariance between individuals calculated from all vertex measurements, which we will refer to as the brain relatedness matrix (BRM). Off diagonal elements of the BRM can be interpreted as a measure of brain similarity between two individuals (see **S2 Appendix**). Finally, 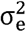 and 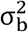 are the variance components for the random effects **e** and **b**. For context, this model is analogous to that used in complex trait genetics to estimate SNP-based heritability, where a Genetic Relatedness Matrix (GRM) replaces the BRM (Yang et al., 2010; Yang et al., 2011). The element i,j of the BRM can be calculated as the inner product of brain measurements of individuals i and j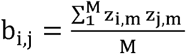. Here, *Z*_*i,m*_ represents the value of vertex m for individual i centred and standardised by its standard deviation over all individuals, z_j,m_ the value of vertex m for individual j centred and standardised over all individuals, M is the total number of vertices or brain features included. We can equivalently use matrix notation, then: 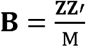, with **Z**_N,M_ a matrix of the centred and standardised brain observations, for N individuals and M brain features. We are interested in estimating the parameters 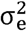 and 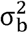 so we can derive the proportion of the trait variance captured by the brain similarities: 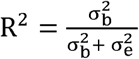. To do so we used the REstricted Maximum Likelihood (REML) method (Patterson and Thompson, 1971) implemented in OSCA.

#### 2.3.2. Mixed model with several random effects

Here, we are dealing with several types of brain measurements: cortical vs. subcortical or thickness vs. surface area for instance. To accommodate the different modalities, we can extend the LMM presented above to jointly estimate the variance accounted for by the different types of measurements: **Y= Xβ** + **b**_1_ + **b**_2_ + **b**_3_ + **b**_4_ +**e** now, with 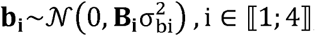, and all other parameters left unchanged. Note that since all **b**_i_ are estimated jointly, each estimate is conditional on the other three parameters fitted in the model. We constructed the BRM **B**_1_ from the cortical thickness measurements, **B**_2_ from the cortical surface area, **B**_3_ from the subcortical radial thickness and **B**_4_ from the subcortical curvature. The variance components 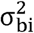 quantify the specific variance attributed to each type of measurement and the quantity 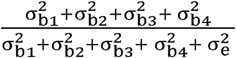 represents the proportion of the trait variance captured by all our brain measurements not biased towards the cortical measurements.

#### 2.3.3. Bivariate models to estimate grey-matter correlation

Finally, we are interested in estimating the correlation (or covariance) between two traits that is attributable to the same brain similarities, which we call grey-matter correlation r_GM_. This can be achieved by fitting a bivariate LMM, a direct extension of the models presented above (Thompson, 1973). We used the AI-REML algorithm in GCTA (Lee et al., 2012) as the multivariate option is not yet available in OSCA. We restricted our bivariate analysis to variables that were significantly associated with grey-matter structure. We derived the residual correlations (r_E_) from the phenotypic (r) and grey-matter correlations estimated by GCTA: 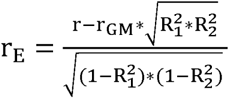 with 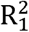 and 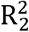 the brain-morphometricity of the two traits included in the bivariate model. For significance testing, we derived SE of r_E_) from a first order Taylor series approximation (delta method, see **Appendix S3** and (Bijma and Bastiaansen, 2014; Lee et al., 2012; Visscher, 1998)).

### 2.4. Covariates used

Our baseline model included commonly used covariates in MRI analyses: acquisition variables (UKB imaging wave, processing with T1w or with combined T1w+T2w), age, sex, and head size (intra-cranial volume (ICV) as well as left and right total cortical surface area and cortical thickness that correspond to the measurements used here). In a follow-up analysis, we included other covariates such as height, weight and BMI to evaluate their confounding effect on the reported associations. We reported the associations between phenotypes and covariates using the adjusted R-squared calculated from linear models estimated in R3.3.3 (R Development Core Team, 2012). As some of the covariates are correlated we report the R^2^ calculated by adding progressively the covariates (same order as above). Thus, the fixed effect R^2^ should not be compared between covariates, but can be contrasted between phenotypes or with the random effect R^2^. We compared the covariates’ associations with our phenotypes of interest in the UKB discovery and replication samples and found highly concordant results between the two samples (**Figure S1**). Thus, any brain-morphometricity difference found between UKB discovery and replication sample should reflect a true difference in the phenotype grey-matter structure association.

### 2.5. Test statistics in mixed linear models

We tested whether the variance accounted for by the brain similarities was significantly different from 0 using a likelihood ratio test on nested models (with and without the random effect). The test statistic follows a chi-square distribution with x degree of freedom (x being the number of variance components tested) for a 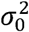 value inside the parameter space. However, when testing 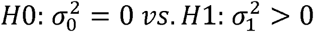, the p-value should be interpreted with caution as the estimator may not be asymptotically normally distributed because 0 is a boundary of the parameter space (Self and Liang, 1987; Stram and Lee, 1994). Some have suggested that the p-value could be better approximated using a mixture of chi-square distributions in the test of significance (Self and Liang, 1987; Stram and Lee, 1994). However, a 50:50 mixture has been shown to be sometimes inappropriate (Crainiceanu and Ruppert, 2004; Pinheiro and Bates, 2000) as the test relies on assumptions often not met in LMM (such as i.i.d. observations) (Crainiceanu and Ruppert, 2004). Thus, we preferred using a χ ^2^ (x df.), the only consequence being a less powerful hence conservative test (Bates et al., 2015; Crainiceanu and Ruppert, 2004; Pinheiro and Bates, 2000). Such test is implemented in OSCA (Zhang et al., 2019), as well as in GCTA (Yang et al., 2011).

### 2.6. Statistical power of the current analyses

In the UKB discovery sample (assuming N=9,500), we have 80% power to detect an effect >2.2% of variance accounted for by the combined BRM (gathering all features), while taking into account multiple testing (pvalue significance threshold p<0.05/175, to ensure a type I error<5%). In the HCP sample (assuming N=1,000), considering the number of tests performed (p<0.05/160), we would need an effect of 20% of variance accounted for to yield the same power (**Appendix S4 and** (Visscher et al., 2014)). For brain correlations, the calculation of statistical power depends on the sample size (set to 9,500), the variance accounted for in each phenotype (we chose 5%), the phenotypic correlation (set to r=0.2), the significance threshold (p<4.2e-5, based on our number of tests) as well as the variance of off-diagonal elements of the BRM *var*(*B*_*ij*_) (0.00096, for the BRM of all brain features) (Visscher et al., 2014). In this example, we had 80% power to detect a brain correlation greater than 0.35, but only a 7% power for a brain correlation of 0.2. Using a sample of N=1,000, as per the HCP, and selecting phenotypes with >20% variance accounted for (everything else being equal), we have a 1% power to detect a brain correlation of 0.35, and we would need a brain correlation greater than 0.99 to achieve 80% power.

### 2.7. Vertex level associations of specific brain features and regions

We conducted post-hoc analyses to identify associations with each type of brain measurement (i.e. left or right measurements of cortical thickness, cortical surface area, subcortical curvature and subcortical thickness) in each cortical (Desikan-Killiany atlas (Desikan et al., 2006)) or subcortical region. For this, we used BRMs specific to each region and brain measurement. Brain regions of interest (ROI) contained between 272 and 12,179 vertices in the left cortex, and between 369 and 11,878 for the right hemisphere. The smallest ROI was the frontal pole and the largest the superior frontal gyrus. Subcortical structures ranged from 930 vertices (Accumbens) to 2502 (Caudate, Hippocampus and Putamen) (Gutman et al., 2013; Gutman et al., 2012). We used the same covariates as in previous LMMs.

### 2.8. In sample prediction (10-fold cross-validation)

We derived brain prediction scores using the Best Linear Unbiased Predictors (BLUP) (Henderson, 1950, 1975; Robinson, 1991) and evaluated them in the UKB discovery sample using a 10-fold cross-validation design. Note that the BLUP predictor was derived from the LMM (REML) analysis described above. When measuring the correlation between grey-matter scores and observed value, we controlled for the same covariates used in the LMMs and included dummy variables to account for hypothetical differences between the groups selected in the cross-validation design. BLUP estimates the predicted values of the random effects (here, **b**, see equation 1) instead of relying on the estimates of fixed effects for all brain features (Goddard et al., 2009; Robinson, 1991). In short, BLUP scores integrate the correlations between vertices to derive weights that correspond to the joint effects of all the vertices. BLUP have desirable statistical properties: they are unbiased and are best predictors in the sense that they minimise the mean square error in the class of linear unbiased predictors (Henderson, 1975; Robinson, 1991), leading to more accurate prediction than other linear predictors (Robinson et al., 2017; Vilhjalmsson et al., 2015). Among others, BLUP scores are routinely used in animal breeding (Robinson, 1991), prediction of individual genetic risk (Robinson et al., 2017) as well as to calculate transcriptomic or methylation age (Peters et al., 2015). BLUP predictors can be calculated in OSCA (Zhang et al., 2019) from summary statistics (analogous to GCTA-SBLUP (Robinson et al., 2017)) and known correlation between vertex measurements.

### 2.9. Out of sample prediction

Finally, we derived BLUP brain prediction scores constructed from the UKB discovery sample, and applied them to the UKB replication and HCP participants. We evaluated the predictive performance using the correlation between grey-matter score and corresponding observed phenotype, controlling for covariates used in the LMMs.

### 2.10. Application of LMMs to identify “best” cortical processing

Here, we defined as “best” processing the MRI cortical processing that maximises the association with a trait of interest, from the minimal number of features (vertices). Thus, we evaluated which of our 10 FreeSurfer processing (fsaverage – no smoothing; fsaverage – smoothing fwhm5, 10, 15, 20, 25; fsaverage6, 5, 4, 3 – no smoothing; ENIGMA ROI processing; **see 2.1.3**) maximised the brain-morphometricity, for all UKB traits considered.

As the ENIGMA processing only consists of 150 measurements (14 subcortical volumes measurements, cortical surface or thickness averaged over 78 ROI defined by the Desikan-Killiany atlas (Desikan et al., 2006)) we used generalised linear models (GLMs – multiple regression) to estimate the brain-morphomometricity. For context, the LMM approach used in the vertex level analyses is a direct extension of GLMs that allows the number of features to exceed the number of participants (p>N).

### 2.11. Data and code availability statement

Data used in this manuscript is held and distributed by the HCP and UKB teams. We have released the scripts used in image processing and LMM analyses to facilitate replication and dissemination of the results (see **URLs**). We have also released BLUP weights to allow meta-analyses or application of the grey-matter scores in independent cohorts.

## 3. Results

### 3.1. Associations between phenotypes and all grey-matter structure vertices

For the phenotypes of interest, we summarised in circular barplots (**Figure 1**) the association (R^2^) with all 654,386 vertex-wise grey-matter measures extracted during image processing, as well as with covariates (acquisition, age, sex and brain/head size variables - see **Methods**). The R^2^ may be interpreted as the proportion of variance in a phenotype captured by all grey-matter morphology. **Figure 1** shows only the significant results (Bonferroni significance threshold; p_UKB_discovery_=0.05/175=2.8e-4, p_HCP_=0.05/160=2.9e-4) with the full results available in **Dataset S3, S4** (see **Figure S2** for positive control associations).

**Figure 1:**
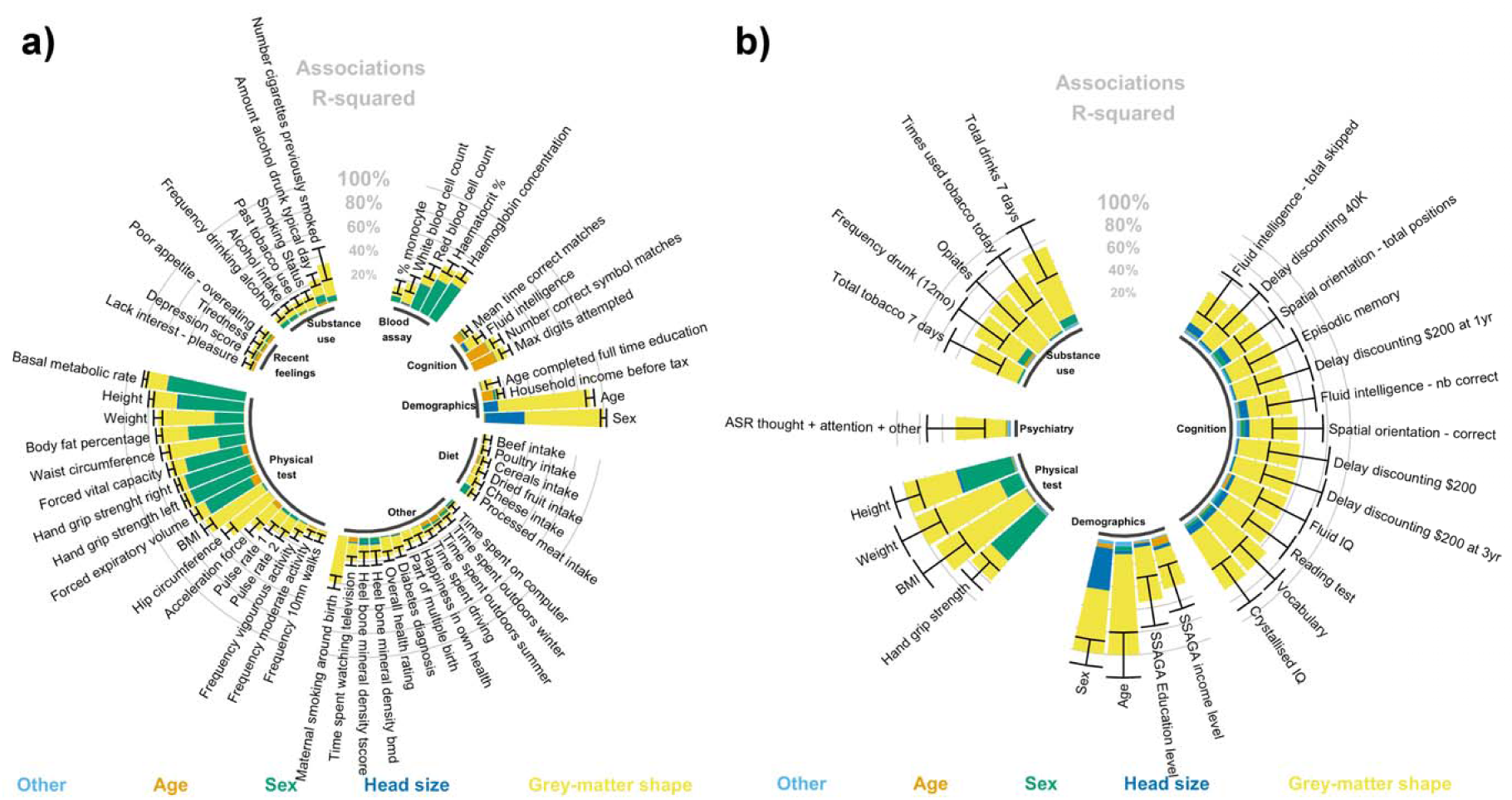
Circular barplot of the associations (R^2^) between phenotypes and grey-matter structure vertices (morphometricity) For clarity, we only plotted the significant associations in the UKB discovery (panel **a**) and HCP sample (panel **b**). We applied Bonferroni 442 correcting to account for multiple testing in each sample. The black bars represent the 95% confidence intervals of the morphometricity 443 estimates. For context, we also present the association R^2^ between phenotypes and covariates of the baseline model, as per the legend under 444 the barplot. As some covariates may be correlated, the R^2^ was calculated by adding progressively the covariates in that order: acquisition and 445 processing variables (labelled “other”), age, sex and head size (ICV, total cortical thickness and surface area). Age and sex were not included as covariates when studying them as phenotypes. See **Dataset S3-4** for full results. See **Figure S1** for positive control associations.

Grey-matter structure was strongly associated with age (R^2^ _UKB_ =0.77, SE=0.018; R^2^ _HCP_ =0.88, SE=0.10), sex (R^2^ _UKB_ =0.66, SE=0.012; R^2^ _HCP_ =0.56, SE=0.059), as well as height (R^2^ _UKB_ =0.22, SE=0.011; R^2^ _HCP_ =0.47, SE=0.060) weight (R^2^ _UKB_ =0.47, SE=0.019; R^2^ _HCP_ =0.81, SE=0.099) and BMI (R^2^ _UKB_ =0.57, SE=0.024; R^2^ _HCP_ =0.92, SE=0.12). Measures of build, body fat and metabolism were also associated with grey-matter structure (R^2^ _UKB_ =0.45, SE=0.019 with waist circumference, R^2^ _UKB_ =0.24, SE=0.013 with body fat percentage, R^2^ _UKB_ =0.19, SE=0.009 with basal metabolic rate; corresponding measures not available in the HCP dataset). In addition, grey-matter structure was associated with measures of strength in both samples (e.g. hand grip: R^2^ _UKB_ =0.074, SE=0.009; R^2^ _HCP_ =0.23, SE=0.58) and levels of physical activity (R^2^ _UKB_ ranging between 0.059-0.25, not-significant in the HCP).

Grey-matter structure was further associated with cognitive domains (R^2^ _UKB_ ranging in 0.048-0.13, R^2^ _HCP_ in 0.34-0.57), smoking (R^2^ _UKB_ ranging in 0.11-0.28, R^2^ _HCP_ in 0.45-0.65), alcohol consumption (R^2^ _UKB_ ranging between 0.071-0.14, R^2^ _HCP_ =0.63, SE=0.13), educational attainment (R^2^ _UKB_ =0.097, SE=0.029; R^2^ _HCP_ =0.39, SE=0.11) and income level (R^2^ _UKB_ =0.042, SE=0.014; R^2^_HCP_=0.32, SE=0.10). Associations with diet, blood assay results, depression score and symptoms, diabetes, bone density, lifestyle and maternal smoking around birth were only observed in the UKB, the phenotypes not being available in the HCP (**Figure 1**).

We replicated 23 of the 58 associations listed above in the UKB replication sample (p<0.05/58; **Figure S3a**). Replication of blood assay phenotypes was limited due to the small sample sizes (N∼300), being only collected for the first imaging waves. Beyond statistical significance that depends on sample and effect sizes, the brain-morphometricity estimates were highly similar between the discovery and replication UKB samples (cor=0.95, excluding blood assay, **Figure S3b**). Full replication results have been added to **Dataset S4**.

In the UKB (discovery), results and conclusions did not change regardless of fitting a single random effect or several random effects each corresponding to one of the grey-matter modalities (i.e. cortical thickness, cortical surface, subcortical thickness, and subcortical area) (**Figure S4**). In the HCP, we observed 3 extra significant associations between grey-matter structure and cocaine (urine test), self-reported number of times used cocaine or hallucinogens. Similar to the association found with opiate (urine test), the small number of positive participants warrants replication. We chose not to include these 3 variables in the subsequent analyses.

### 3.2. Controlling for body size

The large associations between grey-matter structure and height, weight, BMI, waist and hip circumference (even after controlling for acquisition, age, sex and head size differences, **Figure 1**) led us to perform a sensitivity analysis to evaluate their contribution to the brain-morphometricity of the traits studied. We repeated the analysis further controlling for height, weight and BMI, which yielded lower R^2^ estimates (**Figure S5**) and fewer significant associations with grey-matter structure. Thus, when correcting for height in the UKB, 4 of the 58 associations with grey-matter structure did not remain significant: household income, monocyte percentage, beef intake, and time spent using computer, see **Dataset S3**). Such finding is consistent with the reported association between body size and income or socio-economic status in the UKB (Tyrrell et al., 2016). When further correcting for weight and BMI another 14 associations did not remain significant including educational attainment, frequency drinking alcohol, most diet items (cereal, dried fruits, poultry, processed meat), time spent driving, red blood cell count, frequency of walks and small exercise. Notably, the brain-morphometricity of the depression score could be completely explained by differences in weight and BMI (R^2^ _baseline_ =0.050, SE=0.018; R^2^ _baseline+height_ =0.048, SE=0.017, R^2^ _baseline+height+BMI+weight_ <0.001, SE=0.007) and none of the associations between grey-matter structure and depression symptoms remained significant (Tiredness, Anhedonia, Poor appetite-overeating, R^2^ _baseline+height+BMI+weight_ <0.014). The brain-morphometricity estimates (correcting for body size) in the UKB replication sample aligned with results from the discovery sample (cor=0.90), except for age and sex showing larger associations with grey-matter structure in the replication analysis (**Figure S6**).

Similarly, 4 of the 27 associations did not remain significant in the HCP dataset: with fluid intelligence (total skipped), hand grip strength, ASR thought and attention problems and frequency of being drunk in the past year. Though we had limited power to detect associations smaller than R^2^ of 0.2 in this sample (see **2.6**).

In light of these results, we chose a conservative approach to present in the main text results that include body size variables as covariates, though the analyses using baseline covariates can be found in the supplementary. We acknowledge (see discussion) that this may be overly conservative, by implicitly making strong assumptions about the direction of causation between body shape, grey-matter morphology and the rest of the phenome. On the other hand, it avoids reporting associations that may be fully or in part caused by differences in body shape.

### 3.3. Grey-matter correlations

We estimated grey-matter and residual correlations (r_GM_ and r_E_) between the phenotypes that showed significant brain-morphometricity in the univariate analyses. r_GM_ can be interpreted as the proportion of grey-matter vertices similarly associated with both traits, while r_E_ offers insight into factors, shared between the traits, but that do not relate to grey-matter structure (e.g. other brain modalities, non-brain contribution). A weighted sum of r_GM_ and r_E_ make up the phenotypic correlation (see **2.3.3**). In this section, we controlled for height, weight and BMI on top of the baseline covariates, which yields a conservative set of 39 phenotypes and prevents results from being confounded by body size (**Figure 2**; **Datasets S5** (UKB), **S6** (HCP) for point estimates). We excluded phenotypes used as covariates (age, sex, head and body size) as regressing them out makes them orthogonal (i.e. not associated) with the remaining traits. We used conservative significance thresholds of 0.05/(35*34)=4.2e-5 for UKB and 0.05/(18*17)=1.6e-4 for HCP that account for the total number of correlations performed in each sample. We highlighted below which grey-matter correlations were also significant in the UKB replication sample (significance threshold p<0.05/ntest i.e. p<1.9e-3).

**Figure 2:**
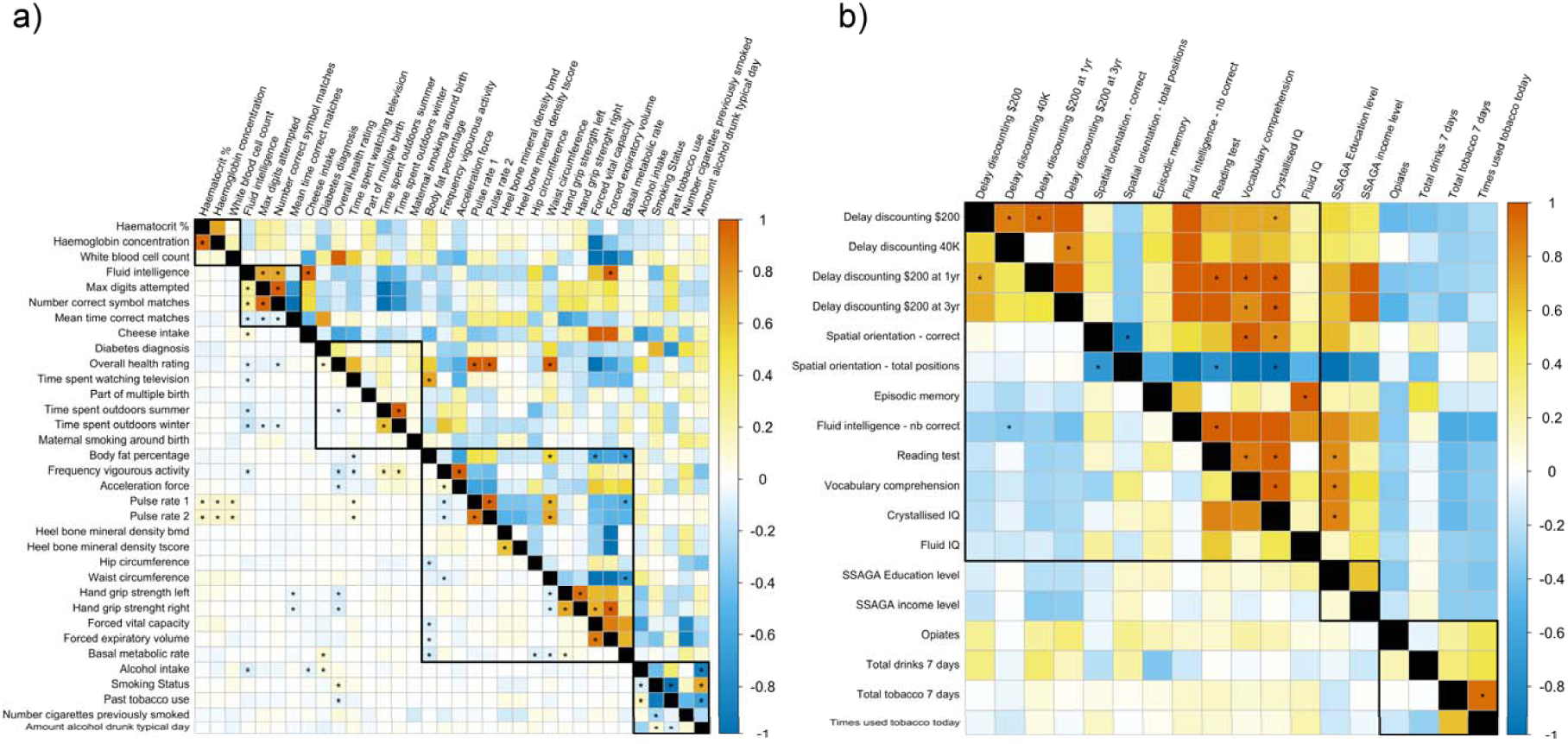
Matrices of grey-matter correlations (upper diagonals) and residual correlations (lower diagonals) between all the variables showing significant morphometricity after controlling for baseline covariates, as well as height, weight and BMI. Panel (a) shows the results for the UKB and panel (b) the HCP results. Correlations significant after multiple testing correction (Bonferroni) are indicated by a star. Blocks circled in black indicate the different phenotype categories used previously (see **Figure 1**). Most grey-matter correlations are observed within categories (e.g. cognition or substance use) but they can also help identifying shared brain-morphometricity between different types of variables (e.g. cheese intake and pulse rate). r_GM_ is a measure of the shared brain-morphometricity between 2 traits and can arise from causal, bi-directional or confounded relationships between phenotypes. Contrasting r_GM_ and residual correlation (r_E_) can indicate how much of the phenotypic correlation is attributable to individual’s resemblance in term of grey-matter structure, compared to other factors (brain or non-brain resemblances).

In the UKB, we observed significant positive grey-matter (and residual) correlations between cognition domains (r_GM_ ranging between 0.71, SE=0.12 and 1.0, SE=0.007; corresponding r_E_ ranging between 0.26, SE=0.014 and 0.94, SE=0.005; **Figure 2**). In addition, we identified grey-matter correlations between measures of physical activity. For example, body fat percentage correlated with waist circumference (r_GM_=0.52, SE=0.12), forced vital capacity (r_GM_=-0.66, SE=0.14), basal metabolic rate (r_GM_=-0.69, SE=0.094, r_GM-replication_=-0.75, SE=0.13) and time spent watching TV (r_GM_=0.73, SE=0.13, r_GM-replication_=0.81, SE=0.16). Pulse rate correlated with waist circumference (r_GM_=0.67, SE=0.13) and basal metabolic rate (r_GM_=-0.55, SE=0.11), acceleration force correlated with frequency of vigorous activity (r_GM_=-0.64, SE=0.17) while hand grip strength (left and right) was associated with forced vital capacity (replicated) and forced expiratory volume (replicated). In addition, we found significant grey matter correlations between substance use phenotypes such as amount of usual alcohol intake and alcohol intake (r_GM_=-0.89, SE=0.086; r_GM-replication_=-1.0, SE=0.12; sign due to coding of the variable, **see Dataset S1**), smoking status (r_GM_=0.71, SE=0.13) and past tobacco use (r_GM_=-0.64, SE=0.14). Finally, we identified unexpected large grey-matter correlations. For example, cheese intake and forced expiratory volume were both correlated (r_GM_=1.0, SE=0.11) with fluid intelligence, and waist circumference correlated with overall health rating and pulse rate (r_GM_>0.67). Overall, 9 out of the 26 significant correlations replicated in the UKB replication sample; sign of the grey-matter correlation was always consistent between discovery and replication analyses (**Table S1**).

In the HCP, we observed positive grey-matter correlations between cognition domains (**Figure 2** and **Dataset S6**) and between IQ dimensions and education level. In addition, the two tobacco related phenotypes were associated with most of the same grey-matter vertices (r_GM_=0.92, SE=0.045). To note, residual correlations and grey-matter correlations were of opposite signs between IQ domains and delay discounting variables, and between cognition and substance use phenotypes. These observations remained after rank-inverse transformation of the variable, suggesting it is not an artefact of the trait distribution. More work is needed to confirm these results in larger samples.

For completeness, grey-matter (and residual) correlations under the baseline model are reported in **Figure S7**, which reveals many large grey-matter correlations between measures of body size and diet, blood assay, activity levels and depression symptoms and score. This further highlights that in the phenome, the brain-morphometricity of some traits may be accounted for by the covariation between these phenotypes and body size measuremements. In particular, depression score was correlated (r_GM_=1) with weight, BMI waist or hip circumference, consistent with its brain-morphometricity lowered to 0 when controlling for body size. In addition, depression score was also correlated (r_GM_=-1) with activity levels and acceleration force, but also with poultry or cheese intake, happiness in own’s health, diabetes and time spent watching television (r_GM_=1), variable themselves strongly associated with measures of body shape (**Figure S7, section 3.2**).

### 3.4. Associations with grey-matter structure of specific cortical and subcortical regions

We investigated the brain-morphometricity of traits by estimating the association with grey-matter structure of specific cortical and subcortical regions (Desikan-Killiany atlas (Desikan et al., 2006)). All phenotypes were corrected for height, weight and BMI in addition to the baseline covariates. Associations with BMI and other body size variables under the baseline model are also presented. In this post-hoc analysis, we used Bonferroni correction to account for the number of tests performed (significance threshold of 0.05/(164*39)=7.2e-6 in the UKB, 1.2e-5 in the HCP).

In the UKB, the largest associations were observed between age of the participants and subcortical volumes (R^2^ ranging between 0.22 and 0.35 for subcortical thickness, 0.20 0.38 for subcortical area), but most cortical regions were also significantly associated with age, albeit to a lesser extent (R^2^ in the 0.0083-0.15 range for cortical thickness, 0.0048-0.15 range for cortical surface area). Next, significant ROI associations included sex, associated with all subcortical volumes (R^2^ in the 0.0049-0.024 range for thickness, 0.0058-0.027 for area) and with many cortical regions (R^2^ in the 0.0011-0.0076 range for cortical thickness, 0.0019-0.014 for cortical surface area) (**Figure S8** and **Dataset S7**). Maternal smoking around birth was further associated with 28 ROI, mostly located in the occipital and temporal lobes (R^2^ in the 0.013-0.026 range with cortical thickness, R^2^ in 0.014-0.071 with cortical surface and R^2^ in the 0.010-0.039 range with subcortical structure). In addition, we found significant associations between cognition domains and structure of thalamus, putamen, pallidum and hippocampus (R^2^ in the 0.0043-0.024 range). Notably, fluid intelligence was associated with all aspects of thalamus anatomy (left and right, thickness and surface area) while the other cognition domains considered were associated with some aspects of thalamus structure. No association between cognition and cortical structure survived multiple testing correction.

Diabetes diagnosis correlated with (left) superior frontal surface area (R^2^=0.054), as well as with thalamus, putamen, and pallidum thickness (R^2^ ranging between 0.0067 and 0.015), or thalamus and hippocampus surface (R^2^ in the 0.0061-0.014 range). Alcohol intake was associated with left thalamus thickness (R^2^=0.018) while smoking status and past tobacco use were associated with thalamus, caudate, putamen and pallidum thickness, as well as with thalamus surface area (R^2^ in the 0.007-0.020 range). Finally, we also observed small associations between cortical or subcortical regions and overall health rating, time spend watching TV, body fat percentage and physiological measurements (**Figure S8**).

Using the replication UKB sample, we replicated 633 out of the 975 significant ROI-trait associations (p<0.05/975). Most associations were found with age, sex and body size variables, though we also replicated associations between subcortical volumes and hand grip strength or time spent watching TV (**Dataset S8**). In addition, the magnitude of the associations with age, and body size were greatly similar between discovery and replication analyses (**Figure S9**). For sex, we observed larger ROI associations in the UKB replication sample (**Figure S9**), consistent with the larger brain-morphometricity observed in this sample (**Figure S6**).

In the HCP sample, age was associated with thickness (R^2^ in the 0.020-0.049 range) and surface area (R^2^ ranging between 0.067-0.10) throughout the cortex, as well as with subcortical structure (R^2^ in the 0.016-0.087 range). Sex was associated with cortical thickness of the lateral orbitofrontal cortex (R^2^ in the 0.059-0.073 range), as well as with subcortical structure (R^2^ in the 0.042-0.19 range). In addition, we found large associations between cocaine, opiate or hallucinogens use and surface area of several cortical regions located in the temporal lobe (fusiform, superior temporal, insula), frontal (pars-triangularis, pars-opercularis, caudal-middle frontal), parietal (supramarginal, superior and inferior-parietal, precuneus) or in the cingulate (R^2^ in the 0.25-1.00 range for cocaine test, R^2^ in the 0.43-0.46 range for opiates, R^2^ in 0.25-0.56 for number of times used hallucinogens). However, the small numbers and possible outliers in the vertex-wise measurements make such associations prone to false positives. Alcohol consumption was also associated with surface are of the frontal cortex (right rostral middle frontal, paracentral and precentral gyri, R^2^ in the 0.28-0.36 range). No other association survived multiple testing correction (**Figure S10** and **Dataset S9**).

Body size variables were strongly associated with subcortical structure under the baseline model (R^2^ ranging between 0.010-0.059 for height, R^2^ between 0.048-0.30 for the others) and to a lesser extent with cortical surface area (R^2^ between 0.0078-0.026 for height, R^2^ between 0.0061-0.060 for the others) and cortical thickness (R^2^ in 0.0039 0.016 for height, R^2^ in 0.0017 0.045 for the others). The associations between grey-matter structure and body size were pervasive (72/164 significant ROIs associations with height, 109 with waist circumference, 105 with BMI) (**Figure S11, Dataset S10**), suggesting that when acting as confounders height, weight or BMI could lead to false positives in many brain regions.

### 3.5. Ten-fold cross-validation in the UKB and prediction into the UKB replication sample

For each UKB participant, we calculated grey-matter scores relative to phenotypes showing significant brain-morphometricity, by estimating the marginal association between each vertex and the trait of interest. As in previous sections, we used height, weight and BMI controlled for baseline covariates; and further regressed out body size for all other phenotypes. We evaluated the prediction accuracy of the grey-matter BLUP scores by computing their correlations with the observed values (10-fold cross validation design).

Most grey-matter scores significantly correlated (positively) with their corresponding phenotypes (significance threshold of 0.05/39=1.2e-3, **Table 1, S3, Figure 3**). Albeit significant, prediction accuracy was overall low (typically r<0.10, including r=0.11 for sex, r<0.09 with cognition, r=0.08 for alcohol intake, r=0.06 with smoking status) except for age (r=0.60), and maternal smoking around birth (r=0.26) whose grey-matter score correlated more strongly with the observed values. We found similar prediction results in the UKB replication sample, with 29 associations reaching significance at p<1.2e-3 (**Table 1, S3**). Prediction accuracy was on par for most traits, though greater in the replication sample for age and sex (**Figure 3, Table 1, S3**), consistent with a larger training sample being used and larger morphomometricity observed in the replication set (**Figure S6**).

**Table 1:** Summary of the prediction accuracy (R^2^) of the BLUP grey-matter scores. We constructed BLUP scores for the 39 UKB variables showing significant morphometricity and evaluated their predictive power in the UKB (10 fold-cross validation) and HCP sample. When the phenotype corresponding to the grey-matter score was not available in the HCP, we chose the closest available (e.g. waist circumference grey-matter score evaluated against BMI). We evaluate the prediction accuracy by fitting GLM controlling for height, weight and BMI as well as for the baseline covariates (acquisition, age, sex and head size); except for (#) denoting associations not controlling for height, weight and BMI. Rows in bold indicate significant association after correcting for multiple testing (p<0.05/39=1.3e-3) both in and out of sample. This reduced table only shows prediction results significant in all 3 scenarios, see **Table S3** for full table of results.

|  | In sample prediction (UKB) |  |  |  | Prediction into UKB replication |  |  |  | Out of sample prediction (HCP) |  |  |  |  |
| --- | --- | --- | --- | --- | --- | --- | --- | --- | --- | --- | --- | --- | --- |
| | r | pvalue | $R^2$ | AUC (SE) | r | pvalue | $R^2$ | AUC (SE) | HCP variable predicted | r | pvalue | $R^2$ | AUC (SE) |
| Age | 0.64 | 0.0e+00 | 0.41 |  | 0.68 | 0.0e+00 | 0.46 |  | Age | 0.15 | 3.1e-08 | 0.024 |  |
| Sex | 0.26 | 0.0e+00 | 0.067 | 0.58<br>(0.0059) | 0.33 | 9.8e-305 | 0.11 | 0.8<br>(0.0064) | Sex | -0.25 | 8.0e-42 | 0.061 | 0.68<br>(0.016) |
| Part of multiple birth | 0.078 | 4.1e-14 | 0.0061 | 0.66<br>(0.022) | 0.13 | 1.5e-03 | 0.016 | 0.72<br>(0.065) | Being a twin | 0.31 | 1.1e-28 | 0.098 | 0.69<br>(0.016) |
| Body fat percentage# | 0.29 | 0.0e+00 | 0.085 |  | 0.31 | 7.7e-190 | 0.095 |  | BMI | 0.21 | 5.6e-13 | 0.045 |  |
| Waist circumference# | 0.39 | 0.0e+00 | 0.16 |  | 0.38 | 2.0e-205 | 0.14 |  | BMI | 0.21 | 3.5e-13 | 0.046 |  |
| BMI# | 0.45 | 0.0e+00 | 0.2 |  | 0.45 | 7.4e-235 | 0.20 |  | BMI | 0.21 | 2.4e-12 | 0.042 |  |
| Hip circumference# | 0.38 | 0.0e+00 | 0.15 |  | 0.36 | 7.3e-143 | 0.13 |  | BMI | 0.21 | 5.2e-13 | 0.045 |  |
| Height# | 0.25 | 6.5e- | 0.062 |  | 0.23 | 2.6e-132 | 0.054 |  | Height | 0.17 | 1.8e- | 0.03 |  |
|  |  | 318 |  |  |  |  |  |  |  |  | 17 |  |  |
| Weight# | 0.39 | 0.0e+00 | 0.15 |  | 0.39 | 5.8e-231 | 0.15 |  | Weight | 0.19 | 1.2e-12 | 0.036 |  |
| Maternal smoking around birth | 0.26 | 9.8e-132 | 0.069 | 0.66 (0.0067) | 0.25 | 1.7e-08 | 0.063 | 0.65 (0.027) | FTND score | 0.19 | 8.9e-04 | 0.037 |  |

**Figure 3:**
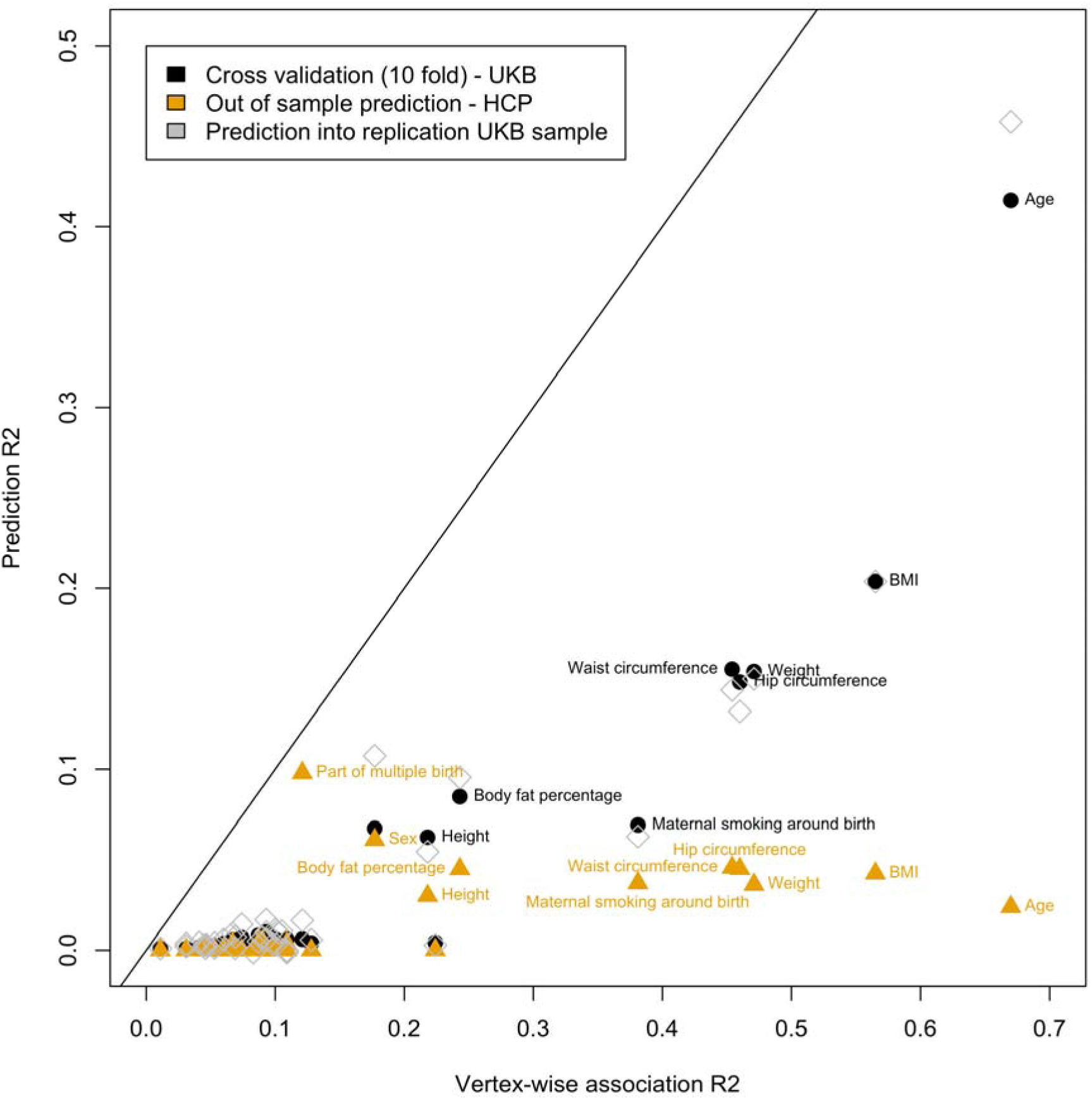
In sample and out of sample prediction accuracy as a function of the total association R^2^. Labels highlight some of the significant prediction having the greatest accuracy. As predicted by the theory, the prediction accuracy is capped by the total association R^2^ (points below the diagonal). In addition, out of sample prediction results in a lower prediction accuracy than in-sample prediction. We hypothesise that the low prediction accuracy of age in the HCP is due to the much younger age range of the HCP participants, compared to the UKB). Participants born from multiple pregnancy appear better identified (predicted) in the HCP than within the UKB sample, which is due to a greater proportion of females and twins in the HCP compared to the UKB, as well as greater morphometricity in the HCP. Such mechanism has been discussed in the field of genetic and solutions exist to correct results for differences in prevalence between samples (Lee et al., 2012). We reported the AUC in **Table 1** (for discrete variables) as it is independent of the proportion of twins and males, thus differences in AUC likely reflect differences in morphometricity between the UKB and HCP samples.

When not correcting for body size, 56/58 BLUP scores significantly correlated with the observed values in the 10-fold cross validation and 42 associations replicated using the UKB replication sample (p<0.05/58, See **FigureS12** and **DatasetS11**). Predicted age correlated with chronological age (r=0.72 in the discovery, r=0.70 in the replication), while predicted sex also strongly associated with the observed value (AUC of 0.90 and 0.89). Grey-matter scores of body shape (under the baseline covariates) were also significantly correlated with the observed values (r=0.25 for height, r=0.29 for body fat percentage, r=0.39 for weight and hip or waist circumference, r=0.45 for BMI). Finally, grey-matter scores of BMI correlated positively with depression symptom count (r=0.10, p-value<1e-14), as expected from the brain-morphometricity of depression being limited the covariation with body size. It even outperformed the grey-matter score built from the depression score itself (r=0.05, p-value<1e.5).

### 3.6. Out of sample prediction – application in the HCP sample

Out of sample prediction validates that the morphometric associations are generalizable to independent brain images, beyond population and scanner differences. We trained our prediction models on the UKB discovery cohort and calculated grey matter scores for each HCP participant. We tested the association between predicted value (brain scores) and the observed phenotype in the HCP. For traits only available in the UKB (e.g. waist circumference) we used a proxy in the HCP (e.g. BMI).

Grey matter scores for age, sex, and being a twin significantly correlated with the observed values (r_age_=0.15, r_sex_=0.25, r_twin-status_=0.31, p-value significant after multiple testing correction) (**Table 1, S3** and **Figure 3**). Grey-matter score for maternal smoking around birth correlated with smoking status (r=0.19). None of the other grey-matter scores significantly correlated with a similar HCP variable.

Without correcting for body size, 19 BLUP scores correlated to corresponding variables (**Dataset S11, Figure S12**). For example, scores for BMI, body fat percentage, hip or waist circumference also correlated positively with BMI (r=0.21, p-value<1.2e-3), while scores for height and weight also correlated with the observed phenotypes (r_Height_=0.17, r_Weight_=0.19). Finally, scores build from diet items or quantifying activity levels significantly predicted BMI in the HCP.

### 3.7. Best cortical processing

We compared the brain-morphometricity estimates obtained by varying the cortical processing options: smoothing of the cortical meshes and applying coarser meshes available in FreeSurfer (see **2.10**). We performed this analysis on the UKB discovery and replication samples as the large SE of the estimates in HCP would limit the interpretation of the results. We used baseline covariates as in **Figure 1**. We found that applying smoothing (5-25mm) or reducing the cortical mesh complexity always led to a lower point estimate of brain morphometricity in the UKB discovery (**Figure 4**) and replication (**SFigure 13, Datasets S12-13 for full tables**) samples. As such, the fsaverage cortical mesh with no smoothing can be deemed “best” processing for all phenotypes considered.

**Figure 4:**
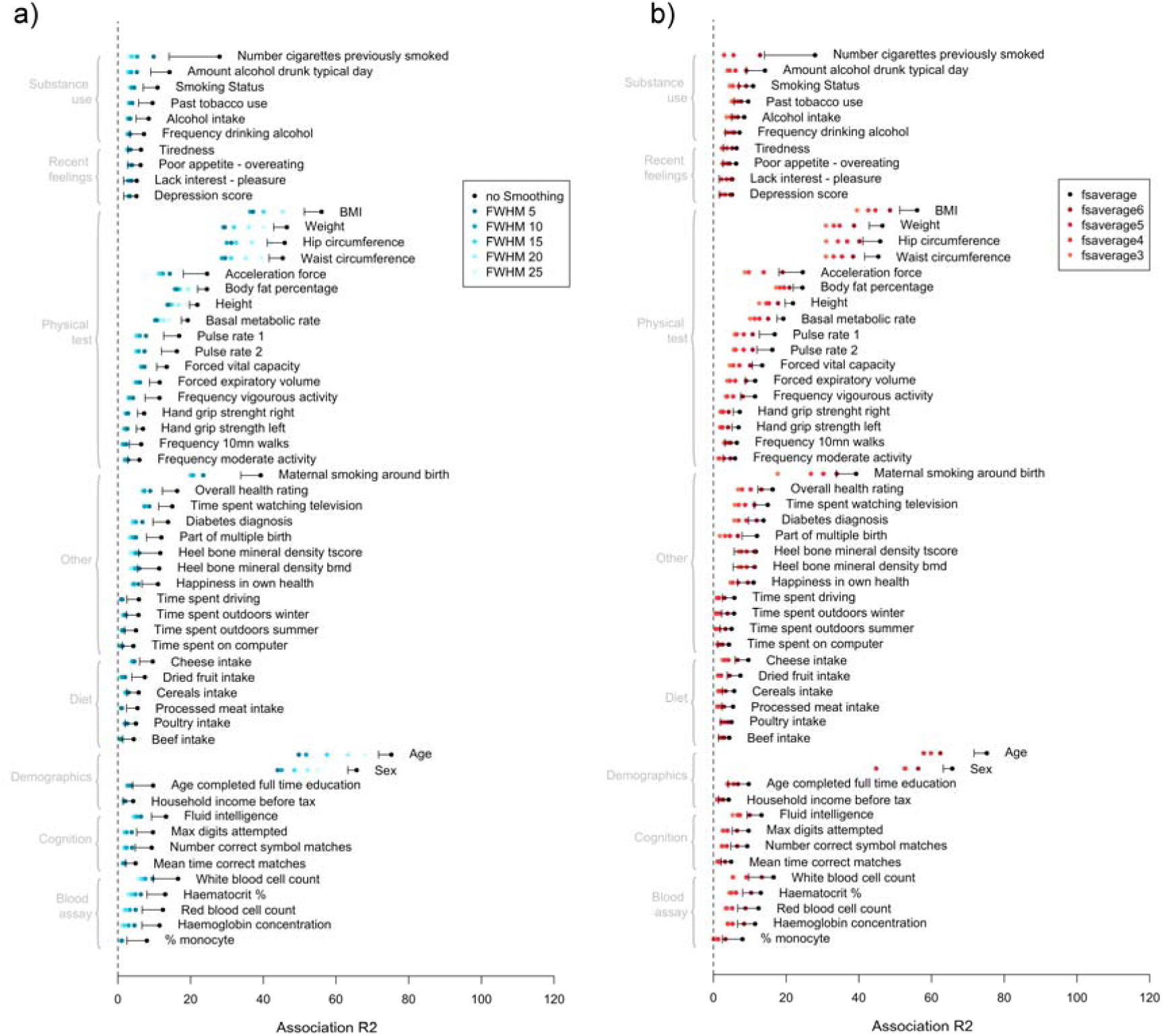
Comparison of brain-morphometricity estimates varying cortical processing options in FreeSurfer. The reduction of brain-morphometricity as a function of mesh smoothing is presented on the left panel (a), while the right panel (b) shows the effect of reducing the cortical mesh complexity. The black bar indicates the lower bound of the 95% confidence interval of the fsaverage-no smoothing estimate (identical to results presented in **Figure1**). Note that brain-morphometricity estimates below the 95%CI lower bound cannot be deemed significantly lower. Rather the 95%CI are presented for context and to remind that all estimate from **Figure 1** do not have the same SE.

In addition, we compared results from **Figure 1** to those from Region-Of-Interest (ROI) based processing (taking the average of each cortical or subcortical region, here ENIGMA processing). We found that the vertex-wise approach always yielded greater association R^2^, thus retained more information than a ROI based dimension reduction (**Figure S14**).

## 4. Discussion

We report the associations between vertex-wise measurements of grey-matter structure and a large set of phenotypes capturing aspects of demographics, physical capacities, substance use, psychiatry, lifestyle and stress/traumas (**Figure 1**). In addition, we introduced the concept of between-trait grey-matter correlation (**Figure 2, Figure S7**) that quantifies the proportion of brain markers shared between two traits. We demonstrated the versatility of our vertex-wise LMM approach by identifying specific cortical and subcortical regions (**Figure S8, S10, S11**) associated with the phenotypes of interest. Finally, we derived BLUP (Best Linear Unbiased Predictor) grey-matter scores and demonstrated their significant predictive abilities in the UKB discovery sample (10-fold cross validation) and in completely independent samples (HCP, and replication UKB, **Table 2, S3, Figure 3**).

Our vertex-wise analyses retained the complexity of the cortical ribbon and subcortical structure, leading to larger associations compared to the standard ROI based data reduction (**Figure S14**). Similarly, reducing the cortical complexity via local averages (smoothing) or halving the number of vertices also led to reduced brain-morphometricity estimates for all phenotypes considered (**Figure 4, S13**). These results indicate that grey-matter scores from “fsaverage-no smoothing” cortical measurements can achieve greater level of prediction, but may require larger training samples to counterbalance their increased complexity (Dudbridge, 2013).

In both the UKB and HCP samples, the largest brain-morphometricity (**Figure 1**) were found between grey-matter structure and age, sex and with measures of body size (height, weight, BMI, body fat percentage, waist or hip circumference). In our post-hoc LMM-ROI analysis, we found those phenotypes to be associated with most cortical and subcortical regions (**Figure S8-11**). Our results for sex are consistent with results from the UKB first release (N=5,216, using ROI average (Ritchie et al., 2018)), while several studies have previously reported associations between BMI and several grey-matter measurements (Cole et al., 2013; Gupta et al., 2015; Kurth et al., 2013; Masouleh et al., 2016; Medic et al., 2016; Opel et al., 2017). Despite such large and widespread pattern of association between body-size and grey-matter we did not observe significant brain-morphometricity for (self-reported) anorexia, bulimia or binge eating though the small numbers (<30 cases in the UKB, **Dataset S1**) limit the interpretability of the results.

We observed moderate to small associations (R^2^<0.4) between grey-matter and substance use (tobacco and alcohol), maternal smoking around birth, blood assay results, education and income level, diet, depression score and symptoms, twin-status as well as cognition domains (**Figure 1**). The latter replicated and expanded the result of an analogous analysis on an early release of the HCP (N=150) and the ADNI dataset (Sabuncu et al., 2016). We note that handedness was only weakly (R^2^_UKB_=0.04, R^2^_HCP_<0.001, not significant) associated with cortical or subcortical grey-matter coherent with the conflicting results reviewed in (Jin Kang et al., 2017). Our results indicate that individuals that display similar grey-matter structure tend to also be similar in term of age, sex, body size, cognition, activity levels, substance use and lifestyle. We did not detect significant association between grey-matter morphometry and psychiatric diagnoses (lifetime self-reported), sleep phenotypes or lifetime stress/traumas (**Dataset S3**) despite previous morphometricity reports from case-control samples of autism, schizophrenia and ADHD (Sabuncu et al., 2016).

When controlling for height, weight and BMI in the analyses, many of the associations became non-significant: such as those between grey-matter and diet, activity levels or depression score/symptoms (R^2^<0.04; **Dataset S3**). Furthermore, we did not detect any significant association between grey-matter structure and other depression related phenotypes (e.g. self-reported diagnosis by a doctor, MDD case-control status as used by the Psychiatric Genetic Consortium (Wray et al., 2018), and neuroticism; **Dataset S3**). Our findings shed a new light on previously published results, as even the largest case-control international initiatives (e.g. ENIGMA-MDD (Schmaal et al., 2016a; Schmaal et al., 2016b)) may reflect, at least in part, variance shared between depression and BMI (such as the causal effect of BMI on depression(Wray et al., 2018)). Understanding the relationship between brain and depression may call to analyse brain regions or features not extracted in the current processing (e.g. brain stem and cerebellum) or features collected from another type of images (e.g. Diffusion Weighted Images (DWI), fMRI).

To summarise, body size is associated with large, widespread variations of grey-matter structure (**Figure 1, Figure S11**) and more work is needed to understand its contribution to published results linking grey-matter anatomy to psychiatric disorders (MDD, bipolar, schizophrenia and substance use are associated with BMI (Luppino et al., 2010; McElroy and Keck, 2012; Rajan and Menon, 2017; Saarni et al., 2009; Wray et al., 2018)) or sexually dimorphic traits (likely associated with height and weight). In addition, body size may be differently associated with the phenome across countries or age groups, which may limit the replication of findings and predictive abilities of body size dependent scores. Note that the possible confounding effects of body size are exacerbated in small case-control samples, leading to increased chances of false positive associations (Button et al., 2013; Ioannidis, 2005). Body size being associated to many brain regions (**Figure S11**), such confounding effect could lead to widespread cortical or subcortical false positives.

In subsequent association and prediction analyses, we made a conservative choice to correct for height, weight and BMI. This meant that we likely reported conservative estimates of brain-morphometricity and fewer significant grey-matter correlations, predictive grey-matter scores or trait-ROI associations (**see Figures S7, S12** and **Dataset S3** for uncorrected results). The large covariation of body-size with the phenome (at least with the variables we selected) is still of interest but it may be more powerful to study directly BMI for example. This is exemplified by the greater prediction accuracy achieved by a BMI grey-matter score (vs. depression specific score) when predicting depression score. Such behaviour can be anticipated based on the large rGM between BMI and depression score (**Figure S7**), combined to the larger brain-morphometricity of BMI (Dudbridge, 2013). Finally, our conservative approach should remind us to be careful when interpreting associations. For example, one should not conclude about actionable links between diet and depression based on the large and significant grey-matter correlations, as it might be mediated by body size. Though, directionality of the associations will need to be established to conclude in this case.

We estimated between-trait grey-matter correlation (**Figure 2, Dataset S5, S6**) that quantifies the proportion of brain markers shared between two traits and found significant relationships between cognition domains, between tobacco and alcohol consumption or between measures of fitness. Large grey-matter correlations between seemingly unrelated traits (e.g. fluid IQ and cheese intake [replicated], waist circumference and pulse rate or overall health rating) raise questions about the nature of the relationships between those variables (causality, true positive or confounded association?). Note that rGM would also capture correlated measurement errors between traits, for example due to head motion or other sources of noise in MRI acquisition.

We further characterised the brain-morphometricity by identifying specific cortical and subcortical regions (ROI) associated with our phenotypes (**Figure S8-S11**). In the UKB, smoking status was associated with thickness and surface of the thalamus (left and right), although we also found associations with the caudate and pallidum. Previous studies have reported association between tobacco usage and volume of left thalamus (Gallinat et al., 2006; Gillespie et al., 2018; Hanlon et al., 2016), which might be due to faster age related volume loss in smokers (Durazzo et al., 2017). We did not replicate other cortical or subcortical associations previously reported (Gallinat et al., 2006; Hanlon et al., 2016; Prom-Wormley et al., 2015). Alcohol intake was also associated with left thalamus thickness in the UKB, consistent with the significant grey-matter correlation (**Figure 2**) between the two traits. The thalamus has been implicated in alcohol-related neurological complications (e.g. Korsakoff’s syndrome)(Pitel et al., 2015) but may also be associated with regular alcohol usage (Cardenas et al., 2007; Pitel et al., 2015) or alcohol use disorder (van Holst et al., 2012). Maternal smoking around birth was further associated with the thalamus, putamen, hippocampus and pallidum, as well as temporal and occipital ROIs. In addition, diagnosis of diabetes was associated with area of the left superior-frontal cortex (**Dataset S8, Figure S8**). Nervous system complications of diabetes (sometimes labelled diabetic encephalopathy) are widely accepted (Mijnhout et al., 2006) but little is known about the specific brain regions associated with the condition (Moheet et al., 2015).

Finally, we derived and evaluated BLUP (Best Linear Unbiased Predictor) grey-matter scores for each individual and the 39 phenotypes showing brain-morphometricity in the UKB (after correcting for body size). The prediction accuracy above what is expected by chance confirmed that the traits associations with grey-matter structure are transferable to independent samples and even samples imaged on a different scanner with different demographics (e.g. HCP, **Table 2, S3, Figure 3**). Overall, the prediction accuracy was below a few percent (of variance) except for age, sex, being a twin, maternal smoking at birth and body size measurements (**Table 2, Figure 3**). Grey-matter score for maternal smoking around birth predicted FTND score in the HCP sample suggesting that passive and active smoking may be associated with similar grey-matter morphology. Our ability to predict (in part) the twin status of participants (**Table 2**) suggests that twins’ grey-matter structure may be more similar than average even if the twins are not from the same family.

Other methods allow to derive prediction from a large number of brain features (e.g. penalised regression, or deep learning) though direct comparison with prediction accuracy from previous publications is limited by the use of different samples, MRI scanners, processing options, input data and prediction algorithm. To note, BLUP is computationally efficient as it does not require estimation of hyper-parameters (as in penalised regression). Similar to polygenic risk scores (Dudbridge, 2013), the prediction R^2^ of grey-matter BLUP scores increases with the training sample size and is capped by the association R^2^ with all vertices (Figure 3). Future application of the grey-matter scores include studying correlates of brain age (Cole, 2017; Cole et al., 2017; Liem et al., 2017), body size and substance use, especially in samples where this information was not collected.

To note, most of the results observed in the UKB discovery sample (brain-morphometricity, r_GM_, ROI based associations) replicated in an independent UKB sample (replication). On the other hand, the UKB and HCP samples differed in term of data collected, age range, country of origin, MRI acquisition, processing and participants’ recruitment, which might explain some of the differences in results (brain-morphometricity of cognition for example).

In the UKB, we chose to add the T2 FLAIR (when available) to improve pial reconstruction in the FreeSurfer processing, though the effect of such option and the possible differences with T1w only processing is not well described in the literature. We observed a large difference in total cortical thickness between participants processed either way (**Figure S2**). This warrants further investigation though it is unlikely to have impacted the results presented here. Indeed, our QC step excluded more than 80% of the 400-odd participants processed using T1w only, likely because they showed outlying brains compared to the T1w+T2 FLAIR processing. In addition, availability of T2 FLAIR was not associated with any of the phenotypes. Finally, the replication of the UKB associations and the out of sample prediction suggest that our results are robust to the presence of these few outliers.

The HCP comprises many twin pairs (thus, non-independent observations), though we modelled the grey-matter relatedness in all analyses, which should account for the grey-matter resemblances arising from shared genetics or environment. A bias due to twins is unlikely as our results on the full HCP sample yielded always similar (e.g. Fluid IQ) or lower (e.g. attention) brain-morphometricity estimates than reported by Sabuncu et al., who selected 1 subject per family (Sabuncu et al., 2016). Finally, the grey-matter similarity of twins was greater than average but in line with the similarity seen between unrelated individuals (**Appendix S2**), which led us not to exclude twin pairs from the analyses (contrary to what is seen/done in genetics).

Due to recruitment choices, the UKB and HCP samples do not contain many psychiatric cases (outside of the highly prevalent MDD, see (Fry et al., 2017) on the healthy volunteer bias in the UKB) and cannot replace the large case-control initiatives (e.g. ENIGMA disease groups). Despite using some of the largest imaging samples available, our ROI based and grey-matter score analyses suffered from limited statistical power, though more data is currently being collected by the UKB.

To complement our analyses (limited to young and older adults), more work is required to understand the relationship between grey-matter morphology and the phenome during development (e.g. in children or adolescents) as well as in specific age/disease groups (Rosenberg et al., 2018), or using different scanners or processing options (e.g. 1.5 Tesla MRI, scanning time, FSL or SPM processing (Flandin and Friston, 2008; Jenkinson et al., 2012)). Note that all associations reported here must be interpreted carefully as they may be causes or consequences of the disorder or trait, or a result of the pervasive pleiotropy underlying human complex phenotypes (Solovieff et al., 2013). Future application or LMM include determining the best MRI image processing for a trait (i.e. the processing options that maximise the association R^2^ ; e.g. **Figure 4**) by extending our analysis to other measures of grey-matter structure (e.g. voxel-based morphometry (Wright et al., 1995)).

We have released the scripts used in image processing and LMM analyses to facilitate replication and dissemination of the results (see **URLs**). We have also released BLUP weights to allow meta-analyses or application of the grey-matter scores in independent cohorts.

## 5. URLs

Summary-level data (BLUP weights) and vertex membership in the Desikan atlas: http://cnsgenomics.com/data.html; OSCA: http://cnsgenomics.com/software/osca/; ENIGMA protocols: http://enigma.ini.usc.edu/protocols/imaging-protocols/;

## 6. Author contributions

PMV, NRW, JY and BCD designed the analyses. FZ and JY developed the OSCA software. YH and BCD created the plots. KK, LY and ZZ assisted BCD with the UKB phenotypic and genetic data, including download, formatting and curation. LS downloaded and processed the HCP MRI images under MJW supervision. BCD downloaded and processed the UKB MRI images. BCD performed the analyses and wrote the manuscript. All the authors reviewed the manuscript.

## 7. Acknowledgements

This research was supported by the Australian National Health and Medical Research Council (1078037, 1078901, 1113400, 1161356 and 1107258), the Australian Research Council (FT180100186 and FL180100072), and the Sylvia & Charles Viertel Charitable Foundation.

Informed consent was obtained from all UK Biobank participants. Procedures are controlled by a dedicated Ethics and Guidance Council (http://www.ukbiobank.ac.uk/ethics), with the Ethics and Governance Framework available at http://www.ukbiobank.ac.uk/wp-content/uploads/2011/05/EGF20082.pdf. IRB approval was also obtained from the North West Multi-centre Research Ethics Committee. This research has been conducted using the UK Biobank Resource under Application Number 12505.

Informed consent was obtained from all HCP participants. HCP Data were provided by the Human Connectome Project, WU-Minn Consortium (Principal Investigators: David Van Essen and Kamil Ugurbil; 1U54MH091657) funded by the 16 NIH Institutes and Centres that support the NIH Blueprint for Neuroscience Research; and by the McDonnell Centre for Systems Neuroscience at Washington University.

We used R(R Development Core Team, 2012) (v3.3.3) for analyses not performed using OSCA (Zhang et al., 2019) and for plots. We used the colour-blind friendly R palette (http://jfly.iam.u-tokyo.ac.jp/color/), *qqman*(Turner, 2014) for QQ-plots, *ggplot2*(Wickham, 2009) and *ggsignif*(Ahlmann-Eltze, 2017) for circular bar plots, *corrplot*(*Wei and Simko, 2017*) for correlation matrix plots, *ukbtools*(*Hanscombe, 2017*) to facilitate UKB phenotype manipulation. Other packages used to assist analyses and data handling include *FactoMineR*(Husson et al., 2015; Husson et al., 2009), *Hmisc(Harrell, 2017), rowr(Varrichio, 2016), pwr(Champely, 2017), XML(Temple and the CRAN Team, 2017), tidyverse(Wickham, 2017a), dplyr(Wickham and Francois, 2015), readr(Wickham, 2017b), reshape2(Wickham, 2007) and rmarkdown(Allaire, 2018)*.

We would like to thank Allan McRae, the Institute of Molecular Bioscience (IMB) and the Research Computing Centre (RCC) IT teams at the University of Queensland for their support with high performance computing, data handling, storage and processing.

