## Supplementary Material for "Widespread associations between grey matter structure and the human phenome"

Supplementary Information for: Widespread associations between grey-matter shape and the human phenome

**This PDF file includes:**

Supplementary text Appendix S1 to S4

Figs. S1 to S14

Tables S1 to S3

Captions for databases S1 to S13

References for SI reference citations

**Other supplementary materials for this manuscript include the following:**

Datasets S1 to S13

Supplementary Information Text

Appendix S1: Evaluation of QC

### Automated quality control based on the BRM

The standards in imaging are to perform a visual QC of the processed images following a (mostly) automated pipeline. For example, the ENIGMA protocol recommends checking participants with outlying measurements but also requires a visual QC of each scan to control the cortical and subcortical parcellation (<http://enigma.ini.usc.edu/protocols/imaging-protocols/)>. This may prove extremely time consuming, especially on large samples such as the UKB that were not available when the ENIGMA pipeline was created.

Here, we propose to utilise the information contained in the BRMs to perform QC. We excluded participants showing extreme values on the diagonals of the BRMs (diagonal>2.5, we did not observe any heavy left tails as per **Appendix S2**). In addition, we excluded individuals showing an outlying level of covariance with other participants, as they could confound our variance component analyses. We took the average of the BRM elements (in absolute value) for each individual (i.e. average of ith row of the BRM for the ith individual) and excluded participants with a statistic more that 4SD away from the mean. We reported the histograms of BRM diagonals and off-diagonals before and after QC (**Appendix S2**). The arbitrary cut-off for the BRM diagonal (>2.5) was determined from the HCP sample on which we had performed visual QC as per ENIGMA protocols (<http://enigma.ini.usc.edu/protocols/imaging-protocols/>).

We applied the same level of QC in the UKB sample but could not compare our approach to visual QC exclusion due to the size of the dataset. Instead, we describe the participants excluded due to failed processing or QC, to check if their exclusion may impact the results presented (**Appendix S2**).

For the 8 BRMs in each sample (left and right hemispheres as well as the different modalities: cortical thickness, cortical surface area, subcortical thickness and subcortical curvature), quality control exclusion removed the heavy tails in the distributions of the diagonal and off-diagonal elements of the BRMs (**Appendix S2**). This suggests that our QC approach should prevent us from false positives caused by outlying values in the BRMs, when estimating the variance accounted for by brain similarities.

### Comparison of visual vs. BRM-based QC approaches in the HCP sample

A total of twenty-four participants were excluded in our QC step based on the diagonal values (>2.5) or extreme level of covariance any of the 8 BRM constructed. Twenty-two participants were flagged using each of the BRM QC criteria. More importantly, 20 outlying individuals were flagged by both BRM criteria. Thus, participants with outlying brains, as indicated by large BRM diagonal elements tended to exhibit outlying covariances with other individuals, potentially causing unstable estimates in variance component analyses.

Out of the 24 individuals excluded in our data driven QC, 14 had also been flagged using the ENIGMA visual QC protocol: 3 were fully excluded for incorrect cortical reconstruction, 7 had an incorrectly segmented hippocampus and 7 others failed visual QC for 3+ cortical regions. Finally, our data driven QC did not identify some individuals flagged using the ENIGMA visual QC: 4 with incorrect hippocampal reconstruction and 108 with incorrect parcellation of the cingulate cortex. The case of the cingulate parcellation is highlighted in the ENIGMA QC protocols as its boundary with regions in the frontal cortex are often misplaced in FreeSurfer. However, this should not be a problem when working at a vertex level as the cortical ribbon remains well segmented, and this may be why these individuals are not identified by our QC approach.

### Description of excluded participants in the UKB

We report the mean (SD) or % of each answer (for qualitative variables) for all the phenotypes considered from the UKB (**Dataset S1**) and compare the mean and variances between included and excluded participants. We used a conservative Bonferroni significance threshold of 1e-4 to account for the number of tests.

The participants we excluded (either for unusable T1, or QC) were on average more than 2.7 years older than the individuals used in the analysis (p-value<3.3e-7) and men were over-represented (62% of excluded were men vs. 47%, p-value<1.8e-5). In addition, excluded individuals were more variable in term of digit matching reaction time, dried fruit intake and exposure to passive smoking. They were less variable than included participants in regard to their basophil percentages (**Dataset S1)**.

Individuals with unusable T1 reported a smaller amount of passive smoking at home (0.001 days a week vs. 0.2, p-value=4.7e-9). They were also less variable than individuals included in the analysis in term of their depression scores.

On the other hand, individuals excluded from QC were 10% less performant as the digit matching task than included participants (smaller number of correct matches or attempted matches, greater reaction time, p-value<4.8e-7). They also were more likely to be diabetics (10% in QCed participants vs. 5%, p-value=1.e-5) and had a reduced acceleration force (-2.4m/s-1, p-value=3.5e-5) as well as greater waist circumference (+3.1cm, p-value=7.0e-7). In addition, the individuals QCed out of the analyses had a greater ICV, smaller grey matter volume, hippocampus volumes or cortical thickness. More importantly excluded individuals exhibited much greater variances in all brain measurements which suggests imperfect/failed processing.

### Sample description HCP

Similar to the results in the UKB, HCP participants excluded by QC showed a significantly greater variance in brain measurements than included participants (**Dataset S2**). This further validates our QC approach, suggesting that the participants QCed out exhibit outlying brains, some due to failure of the MRI processing pipeline.

In addition, excluded participants differed (p-value<1e-4) on some aspects of cognition: delay discounting $200 at 5 years (smaller mean and variance), spatial orientation (total positions; greater variance) and sustained attention (longest run non-response; smaller variance), depression scores (smaller mean and variance).

The similarities between excluded participants in the UKB and HCP (e.g. depression scores or cognition) are intriguing. We hypothesise that these phenotypes may be associated with greater level of movement in the scanner leading to lower image quality and failed processing. Us and others previously showed that inattention and hyperactivity are associated with greater movement in resting-state fMRI(1, 2), and a subsequent study in the HCP found multiple factors also associated with motion during rs-fMRI (for example: some cognition domains, antisocial or somatic scores, weight and BMI as well as tobacco use)(3).

Note that when the variance in excluded and included participants differs, the sample participants may not capture the full phenotypic variance and the results of variance component analyses should be interpreted with caution. In other words, we are estimating the proportion of in sample variance accounted for by brain features which may differ from the proportion of total phenotypic variance accounted for in the population.

Appendix S2: BRM interpretation

Diagonals of the BRM consist of the mean square of the participants’ vertex-wise measurements. Since the vertex-wise data are centred, larger diagonal elements reflect a greater proportion of extreme phenotypes, may they be small or large vertex measurements. Thus, we can interpret large diagonal values as “outstanding brains” in term of size/shape or due to failure in processing (e.g. incorrect cortical or subcortical parcellation). On the other hand, small diagonal values indicate small absolute values across the brain measurements (mean centred), hence brains close to the average brain. Across all the samples and BRMs, we observed that the diagonal elements were centred around 1, and skewed to the right (**Appendix** **S2 Figure 1-2**).

Off-diagonal elements of the BRM are the covariances between two individuals’ measurements, thus greater values indicate greater similarities between the pair of participants. Off-diagonal elements are normally distributed with a mean of 0. Their dispersion varies upon the brain modality considered and the degree of correlation between the vertices (**Appendix** **S2 Figure 1-2**).

Participant’s brain similarities are thought to reflect (some of) the participants’ similarities in genetics and environment, in other words we expect part of the variance accounted for by the BRM to be genetic. Indeed, we observed a positive correlation between elements of the BRM and of the Genetic Relatedness Matrix (GRM) in the UKB or the pedigree matrix in the HCP, which suggests that participants more alike genetically also exhibit more similar brains (**Appendix** **S2 Figure 3-4**). The GRM was calculated in GCTA(4) from the hap-map 3 variants further filtered for MAF>0.01, pHWE<10-6 and missingness<0.05 for a total of 1,123,943 variants. The GRM calculation was restricted to participants of European ancestry (N=456,426) defined by a >0.9 posterior probability of belonging to the 1000G reference ancestry cluster. Overall, the pairs of twins and especially monozygotic twins did not exhibit outlying BRM values compared to the unrelated pairs (**Appendix** **S2 Figure 4**), suggesting that they should not confound the results of a variance component analysis. Quite the opposite, they increase the variance of the off-diagonal BRM elements which results in an increased power **Appendix S4**).

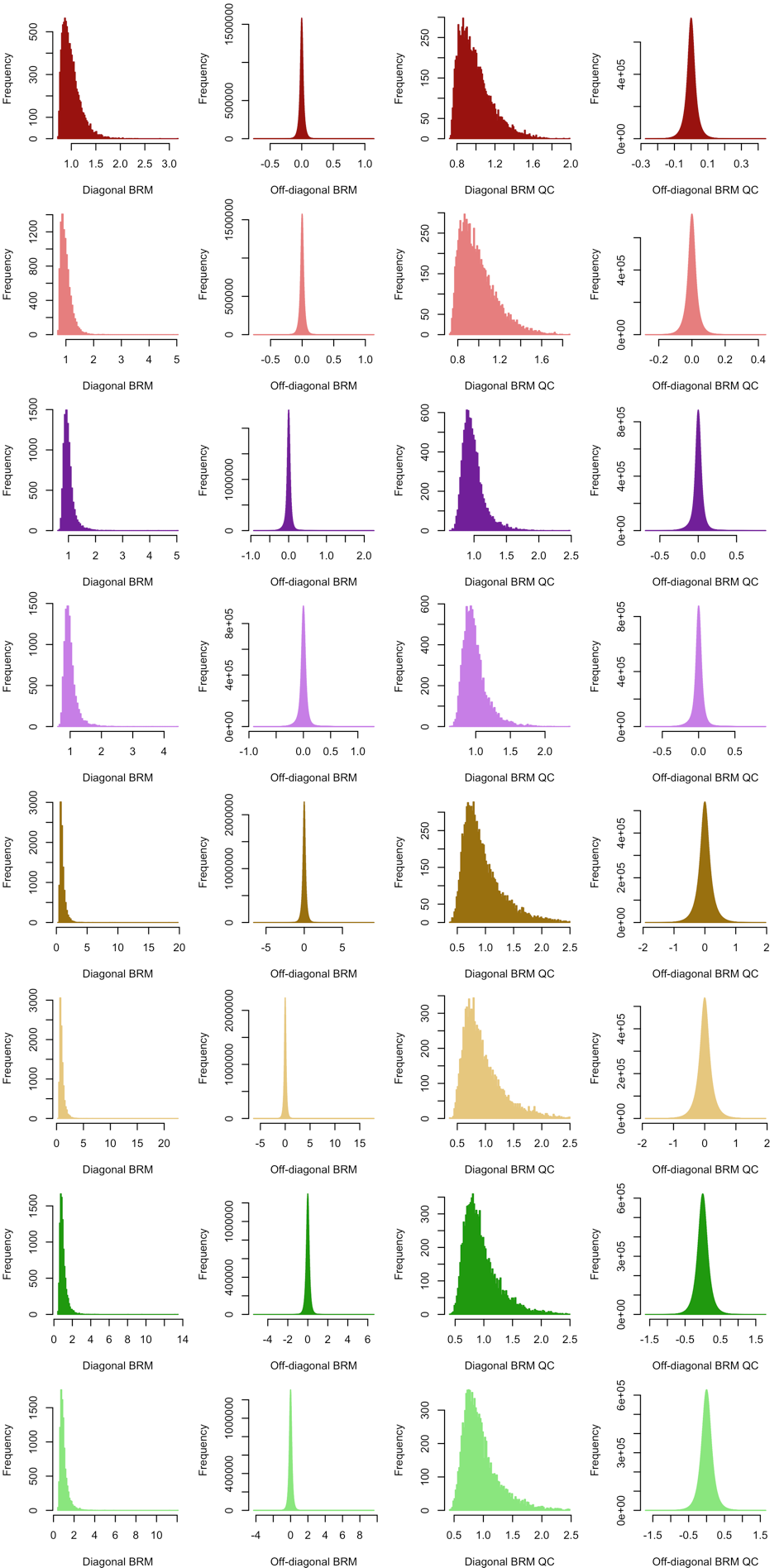

Appendix S2 Figure 1: Histograms of diagonal and off-diagonal elements of the UKB brain relatedness matrix.
Plots are presented before (left panels) and after participants’ QC exclusion (right panels). Colors correspond to the different brain modalities: dark red – left cortical thickness, light red – right cortical thickness, dark purple – left cortical area, light purple – left cortical area, dark yellow – left subcortical curvature, light yellow – right subcortical curvature, dark green – left subcortical thickness, light green – right subcortical thickness.

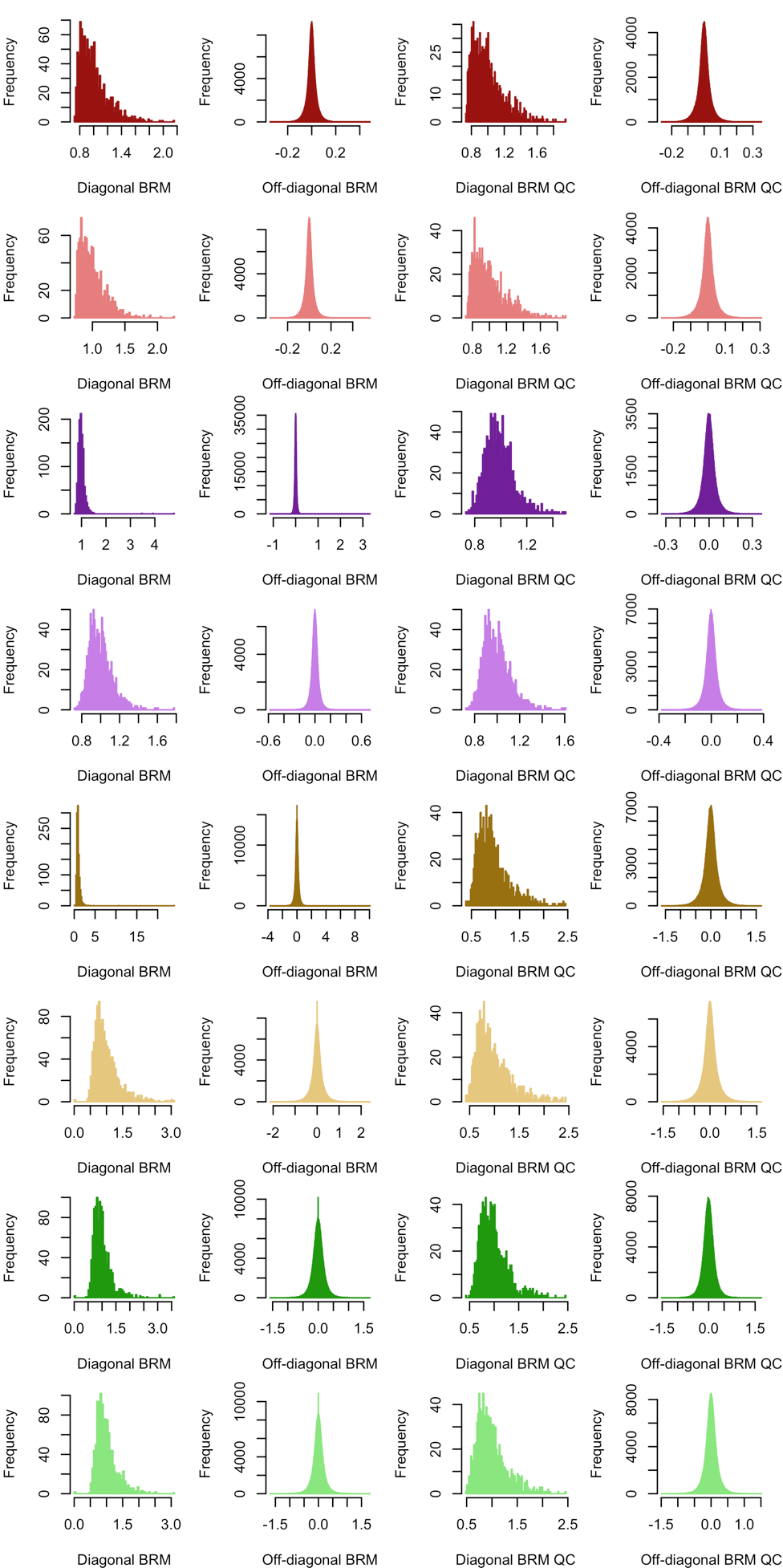

Appendix S2 Figure 2: Histograms of diagonal and off-diagonal elements of the HCP brain relatedness matrix.
Plots are presented before and after participants’ QC exclusion. Colors correspond to the different brain modalities: dark red – left cortical thickness, light red – right cortical thickness, dark purple – left cortical area, light purple – left cortical area, dark yellow – left subcortical curvature, light yellow – right subcortical curvature, dark green – left subcortical thickness, light green – right subcortical thickness.

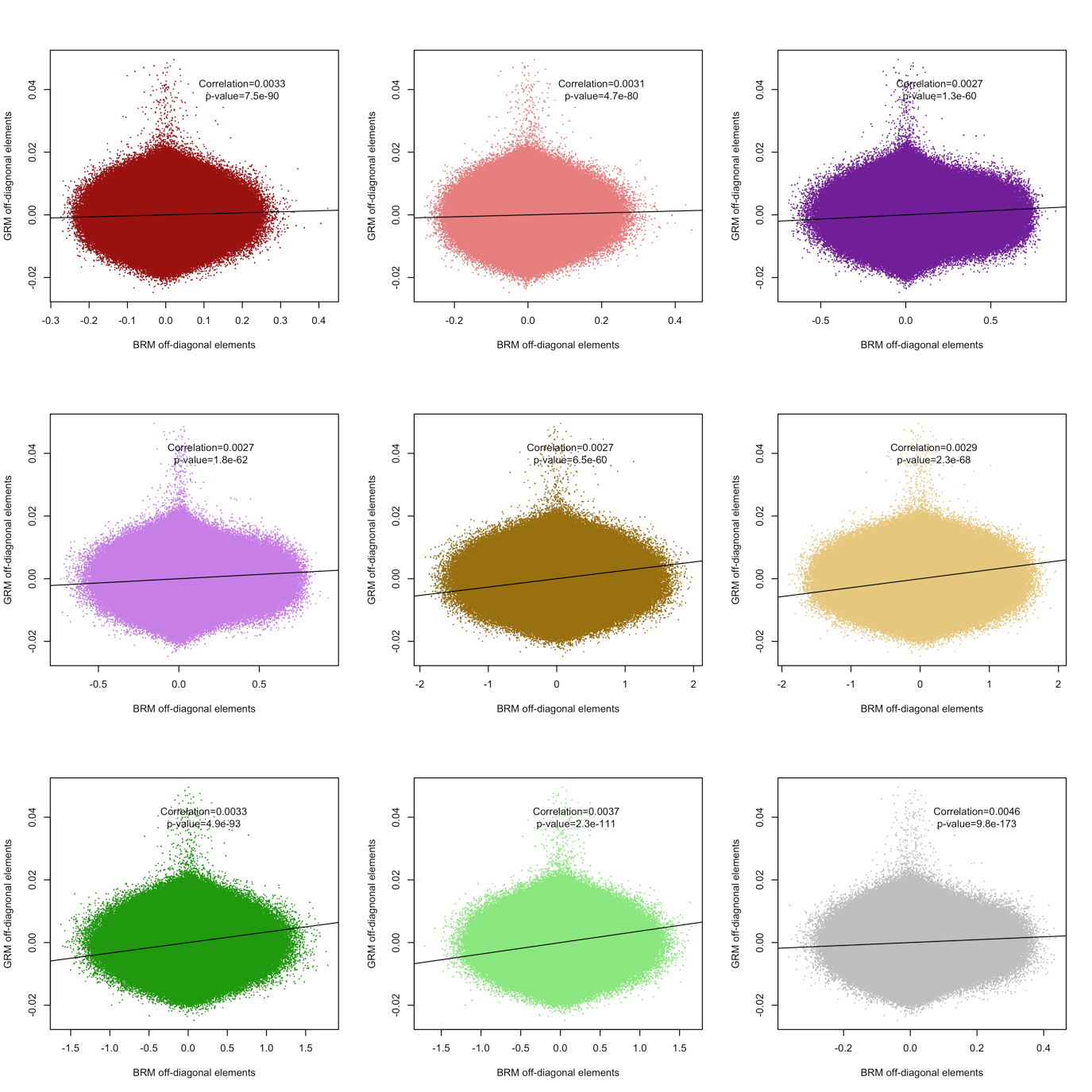

Appendix S2 Figure 3: Scatterplot showing the correlation between GRM and BRM pairwise elements in the UKB sample.
 Correlations and p-values are shown. Colors correspond to the different brain modalities: dark red – left cortical thickness, light red – right cortical thickness, dark purple – left cortical area, light purple – left cortical area, dark yellow – left subcortical curvature, light yellow – right subcortical curvature, dark green – left subcortical thickness, light green – right subcortical thickness. Grey corresponds to all brain vertices.

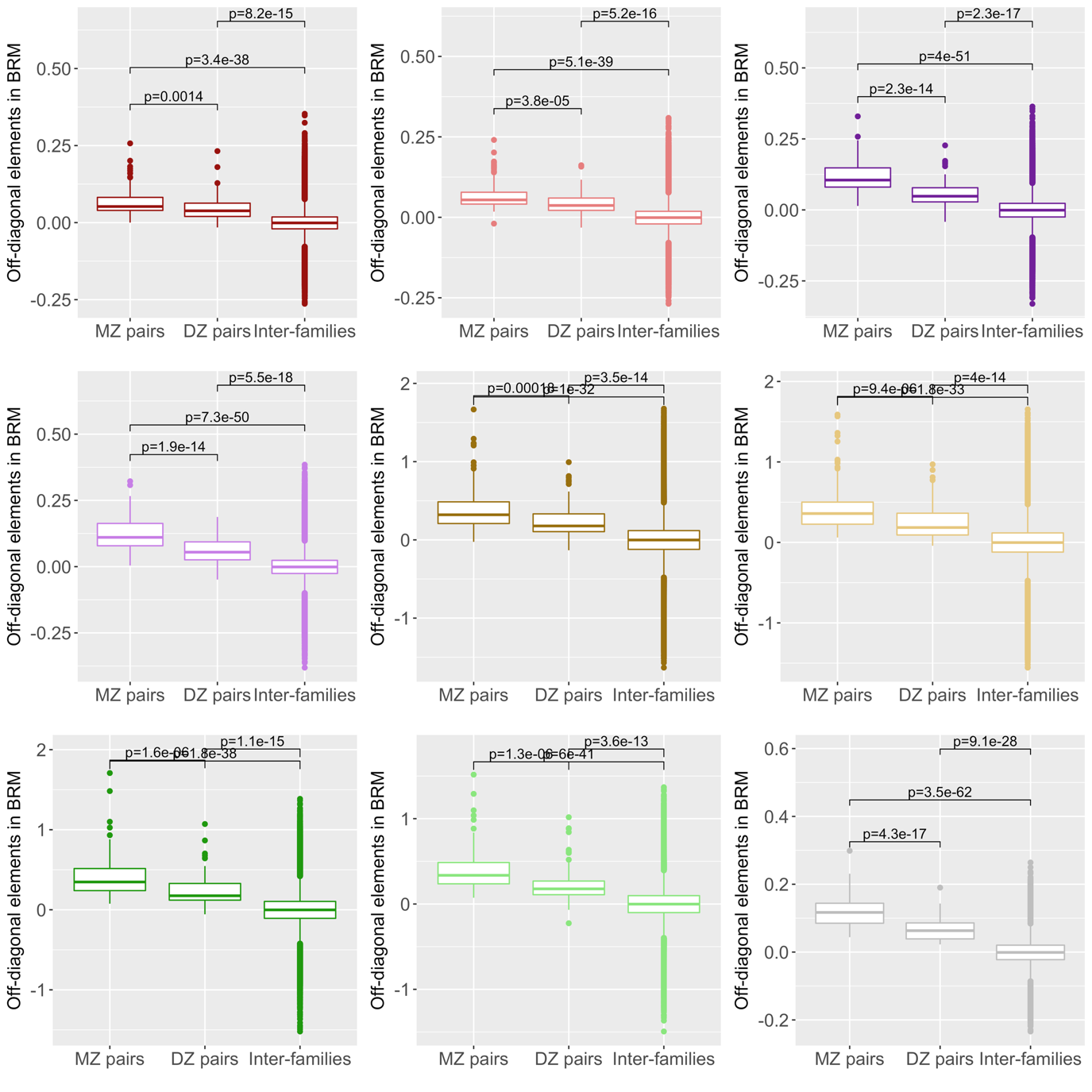

Appendix S2 Figure 4: Boxplot showing the BRM pairwise elements per zygosity group in the HCP.
Tests of mean differences are shown. Colors correspond to the different brain modalities: dark red – left cortical thickness, light red – right cortical thickness, dark purple – left cortical area, light purple – left cortical area, dark yellow – left subcortical curvature, light yellow – right subcortical curvature, dark green – left subcortical thickness, light green – right subcortical thickness. Grey corresponds to all brain vertices.

Appendix S3: SE of the residual correlation

The residual correlation between random variables X and Y can be expressed as a function of the residual covariance between X and Y ($\hat{\sigma}_{XY}$), and the residual association with X or Y (${\hat{\sigma}^{2}}_{X} and {\hat{\sigma}^{2}}_{Y}$).

$$rE=\frac{\hat{\sigma}_{XY}}{\sqrt{{\hat{\sigma}^{2}}_{X}* {\hat{\sigma}^{2}}_{Y}}}=\frac{a}{\sqrt{b*c}}$$

We can derive a first order tri-variate Taylor series approximation around the expected values of $\hat{\sigma}_{XY}, {\hat{\sigma}^{2}}_{X} and {\hat{\sigma}^{2}}_{Y} (denoted \mu_{a}, \mu_{b} and \mu_{c} for convenience)$. Doing so, we implicitly assume these numbers are estimated with a reasonable confidence so that the Taylor series approximation around the mean holds. Thus,

$rE \approx\mu_{rE}+\frac{(a-\mu_{a})}{\sqrt{\mu_{b} * \mu_{c}}}-\frac{0.5 * \mu_{a} * \left( b-\mu_{b} \right)}{\mu_{b} \sqrt{\mu_{b}*\mu_{c}}} -\frac{0.5 * \mu_{a}* (c-\mu_{c})}{\mu_{c} \sqrt{\mu_{c}*\mu_{b}}}$

Taking the variance:

$$V\left( rE \right)\approx\frac{V\left( a \right)}{\mu_{b}* \mu_{c}}+ \frac{0.25*{\mu_{a}}^{2}* V\left( b \right)}{{\mu_{b}}^{3}* \mu_{c}}+ \frac{0.25*{\mu_{a}}^{2}* V\left( c \right)}{{\mu_{c}}^{3}* \mu_{b}}-\frac{\mu_{a}* cov\left( a,b \right)}{{\mu_{b}}^{2}* \mu_{c}} -\frac{\mu_{a}* cov\left( a,c \right)}{{\mu_{c}}^{2}* \mu_{b}}+\frac{0.5 * {\mu_{a}}^{2}*cov(b,c)}{{\mu_{c}}^{2}* {\mu_{b}}^{2}}$$

To conclude, approximating the variance of $rE$ using the formula above, requires the variance components estimates ($\hat{\sigma}_{XY}, {\hat{\sigma}^{2}}_{X} and {\hat{\sigma}^{2}}_{Y}$) from the bivariate model as well as their matrix of sampling variance-covariance for the values ${V(\hat{\sigma}}_{XY}), V\left( {\hat{\sigma}^{2}}_{X} \right), V\left( {\hat{\sigma}^{2}}_{y} \right), cov\left( \hat{\sigma}_{XY},{\hat{\sigma}^{2}}_{X} \right), cov\left( \hat{\sigma}_{XY}, {\hat{\sigma}^{2}}_{y} \right) and cov({\hat{\sigma}^{2}}_{X}, {\hat{\sigma}^{2}}_{y})$.

Such values are estimated in OSCA/GCTA and may be found in the log files outputted during the model fitting. Note that SE of the grey-matter correlation is derived using the same approach. The interested readers may refer to (5-7).

For significance testing, we used a one-sided test based on the test statistic:

$\left( \frac{rE}{SE(rE)} \right)^{2}\sim\chi(1)$

Appendix S4: Power of variance-component analyses

Power calculation of variance component analysis may be derived from the sampling variance of the estimate: $var(\hat{\sigma}_{b}^{2}$), which is the square of the standard error (SE) of the estimate. In REML analyses (e.g. GCTA or OSCA) the SE is estimated from diagonal elements of the inverse of the information matrix, and it has not been derived analytically. Visscher et al.,(8) showed that the SE could be approximated using a simpler model formulation known as Haseman Elston (HE) regression(9), which is the ordinal least square equivalent of the REML approach. As such, HE regression should be slightly less powerful that the REML approach, resulting in a marginal underestimation of the power.

Briefly, HE regression(9) performs a linear regression of the phenotype pairwise product z_ij_=y_i_y_j_ on the off-diagonal elements of the BRM: $B_{ij}$. For a pair of individual i and j:

$$z_{ij}= \mu+b B_{ij}+\varepsilon_{ij}$$

This model includes n=N(N-1)/2 pairwise observations and we can easily show that $b=\sigma_{B}^{2}$ (8, 9). In this simple linear regression framework, assuming that the $\varepsilon_{ij}$ are i.i.d. we can calculate the $var\left( \hat{\sigma}_{B}^{2} \right)=var\left( \hat{b} \right)=\frac{var(\varepsilon_{ij})}{n var(B_{ij})}$.

For centred and standardised phenotypes Y, $var\left( z_{ij} \right)=1$ and therefore $var\left( \varepsilon_{ij} \right)\leq1.$Thus, $var\left( \hat{\sigma}_{B}^{2} \right)\leq\frac{2}{N(N-1) var(B_{ij})}$. As a consequence, the power of variance component analysis may be approximated from the sample size N and the variance of the off-diagonal elements of the BRM. The same formula holds for discrete outcome variables (e.g. sex or disease status) and a similar derivations may be performed for a the bivariate case and the power of brain correlation(8). In our UKB and HCP data, we calculated the variance of the off-diagonal BRM in order to get an approximation of the SE of our variance component estimates. The $var(B_{ij})$ were consistent across sample and across the left and right modalities (**Appendix S4 Table 1**). Such stability of the variance of off-diagonal elements has also been observed for GRM(8, 10). Thus, in our UKB imaging sample, the variance of the off-diagonal GRM was 2.0e-5 (after removing related individuals with GRM elements>0.05), consistent with previous report and analytic derivations from our group(8, 10).

We tried to validate the derivations presented above using simulation. Thus, we simulated 100 normally distributed phenotypes (mean 0, variance 1) and estimated the SE of the variance components using OSCA, varying the sample size (from N=500 to 9,000). We used random subsets of the BRM calculated from the UKB data for the variance-covariance of the random effect. We conducted such analysis for the 8 brain modalities as well as for the global BRM. Results of simulation and approximation theory are presented in **Appendix S4 Table 1, Appendix S4 Figure 1 and 2,** and suggest that the approximation yields realistic, though slightly underestimated, values for the SE of the variance component estimates. Note that HE regression is known to produce underestimated SE at high power as the i.i.d. assumption of errors does not hold anymore(11). Despite being small, the underestimation of SE may lead to overestimate the statistical power using the approximation theory. To circumvent this problem, we provide values for $var(B_{ij})$, derived from our simulation analysis, that result in realistic power calculation using the approximation theory (**Appendix S4 Table 1)**. The statistical power may be calculated (for a hypothesised variance accounted for: $\sigma_{B}^{2}$ and a selected risk α) from the non-centrality parameter of the chi-square statistic $ncp=\left( \frac{\sigma_{B}^{2}}{SE} \right)^{2},$using the R formula 1-pchisq(qchisq(1-α, df), df, ncp) or the GCTA-GREML online power calculator (<http://cnsgenomics.com/shiny/gctaPower/)>.

Results presented below also highlight the greater power of the brain variance component analyses compared to estimation of SNP heritability, as indicated by the smaller SE of the estimates. For example, in a sample of N=1000, we would have >60% power to detect an effect $\sigma_{B}^{2}>0.1$ (with α=0.05), but only a 5.5% power to detect a SNP heritability greater than 0.1 (**Appendix S4 Figure 1 and 2**).

|  |  | Var(Bij) UKB | Var(Bij) HCP | Var(Bij) to use in power calculation (corrected based on our simulations) |
| --- | --- | --- | --- | --- |
| cortical area | Left | 0.0012 | 0.0015 | 0.00047 |
|  | right | 0.0012 | 0.0016 | 0.00046 |
| Cortical thickness | Left | 0.0049 | 0.0021 | 0.0017 |
|  | Right | 0.0057 | 0.0024 | 0.0017 |
| Subcortical curvature | Left | 0.057 | 0.057 | 0.017 |
|  | Right | 0.058 | 0.059 | 0.017 |
| Subcortical thickness | Left | 0.035 | 0.036 | 0.018 |
|  | right | 0.034 | 0.035 | 0.020 |
| All modalities |  | 0.0017 | 0.0014 | 0.00096 |

Appendix S4 Table 1: Variance of off-diagonal elements in the UKB and HCP.

Following our simulation results and to avoid overestimating the statistical power of brain variance-component analysis, we recommend using the values in the right-hand side column in the approximation theory formula (see main test for the formula, or <http://cnsgenomics.com/shiny/gctaPower/> for online power calculator).

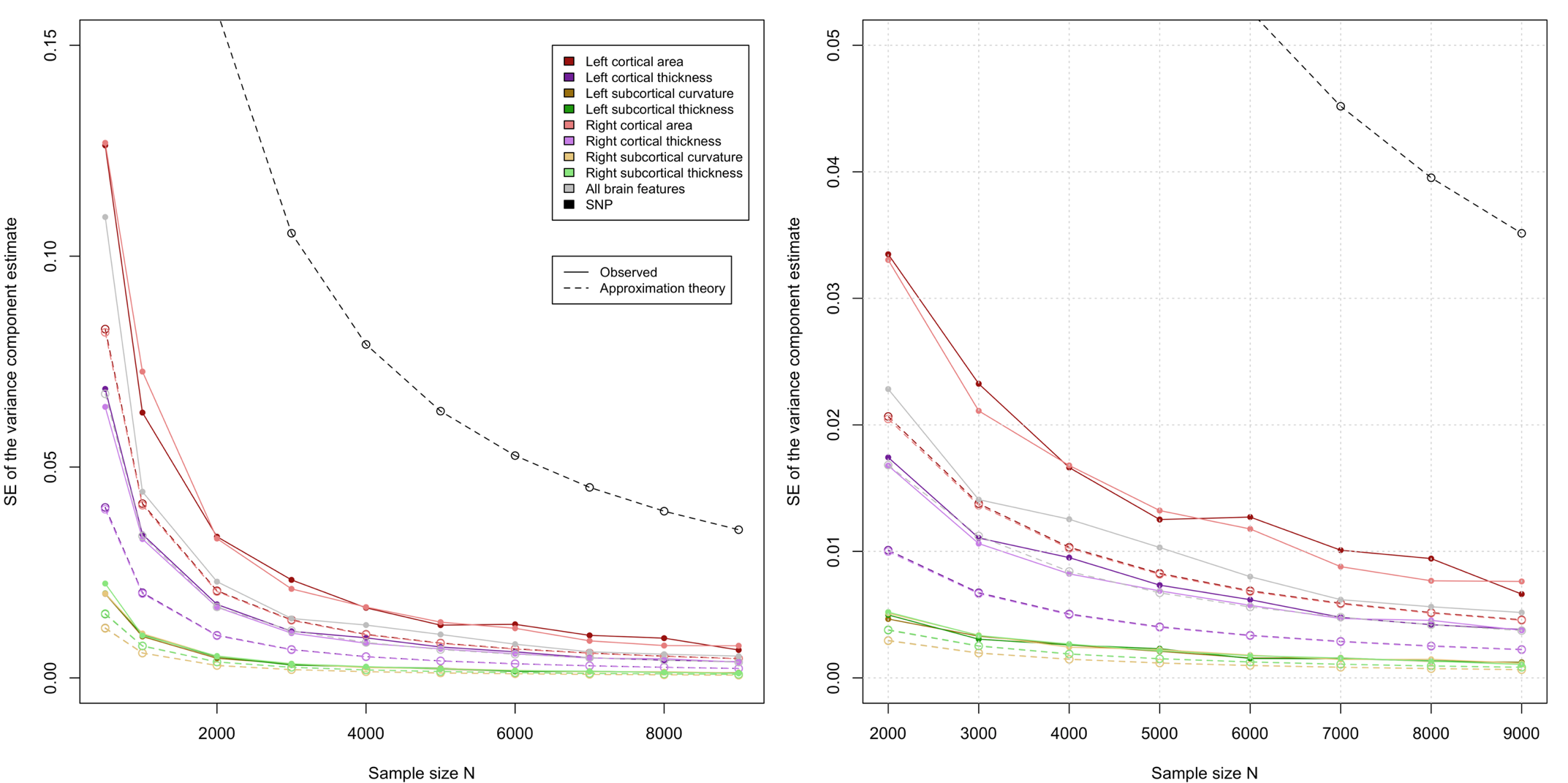

Appendix S4 Figure 1: Empirical and approximated SE of the variance component estimates using BRM and GRM
The right panel is a close up of the left one, restricted to sample sizes above 2000 participants. We did not calculate the empirical power for the genetic case has it has already been shown to match the approximated power (8).

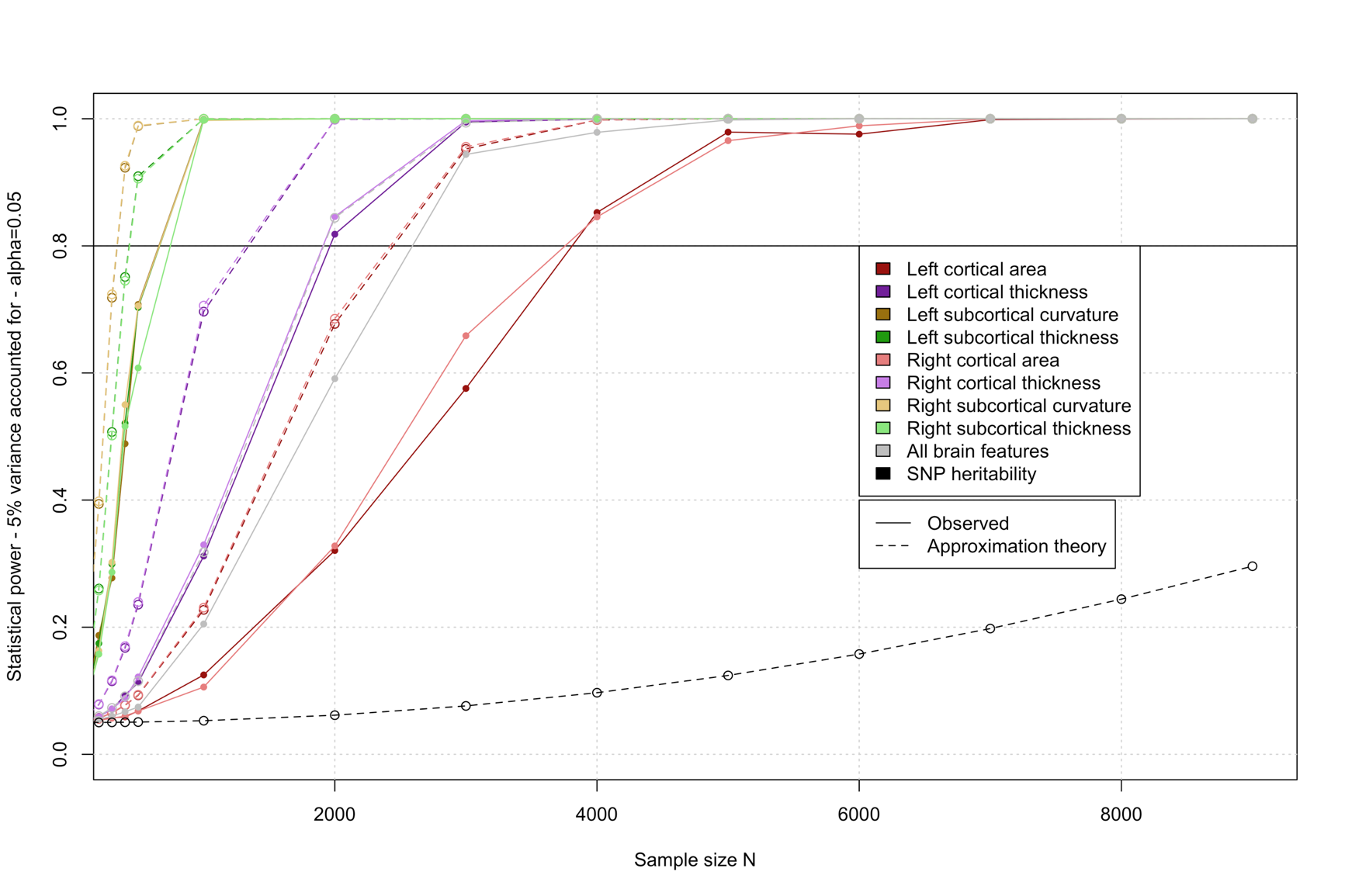

Appendix S4 Figure 2: Statistical power of detecting an association R^2^>5%, with a risk alpha=0.05.
Power derived from simulation and from the approximation theory are compared. We did not calculate the empirical power for the genetic case has it has already been shown to match the approximated power (8).

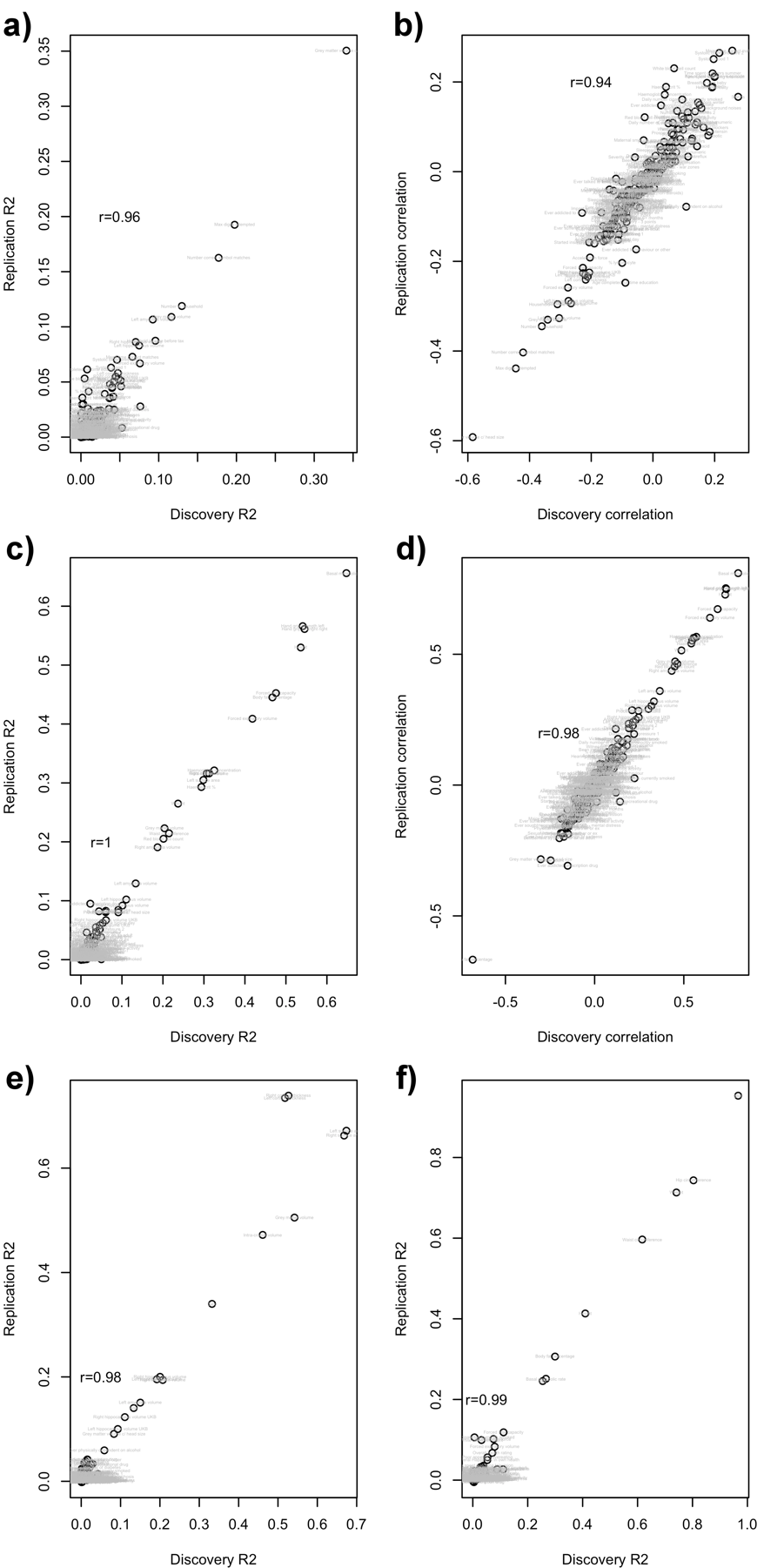

**Fig. S1. Replication R2 (or correlations) are presented as a function of the discovery R2 (or correlation).**

The R2 or correlations correspond to the fixed effect association between a covariate and all other phenotypes (labelled in plots). For example, panel a) shows the association R2 between age and all phenotypes, in the discovery and replication UKB samples. Panel b) shows the same results but using correlations and not R2, thus confirming that the sign was consistent too. Panels, c) and d) show the R2 and correlations between sex and all other variables. Panel e) presents the R2 between phenotypes and head size (ICV, left and right total thickness and area). Panel f) presents the R2 between phenotypes and body size (height, weight and BMI). As head size or body size are composed of several variables, only the R2 is presented. The correlation between discovery and replication results in shown in each panel as r.

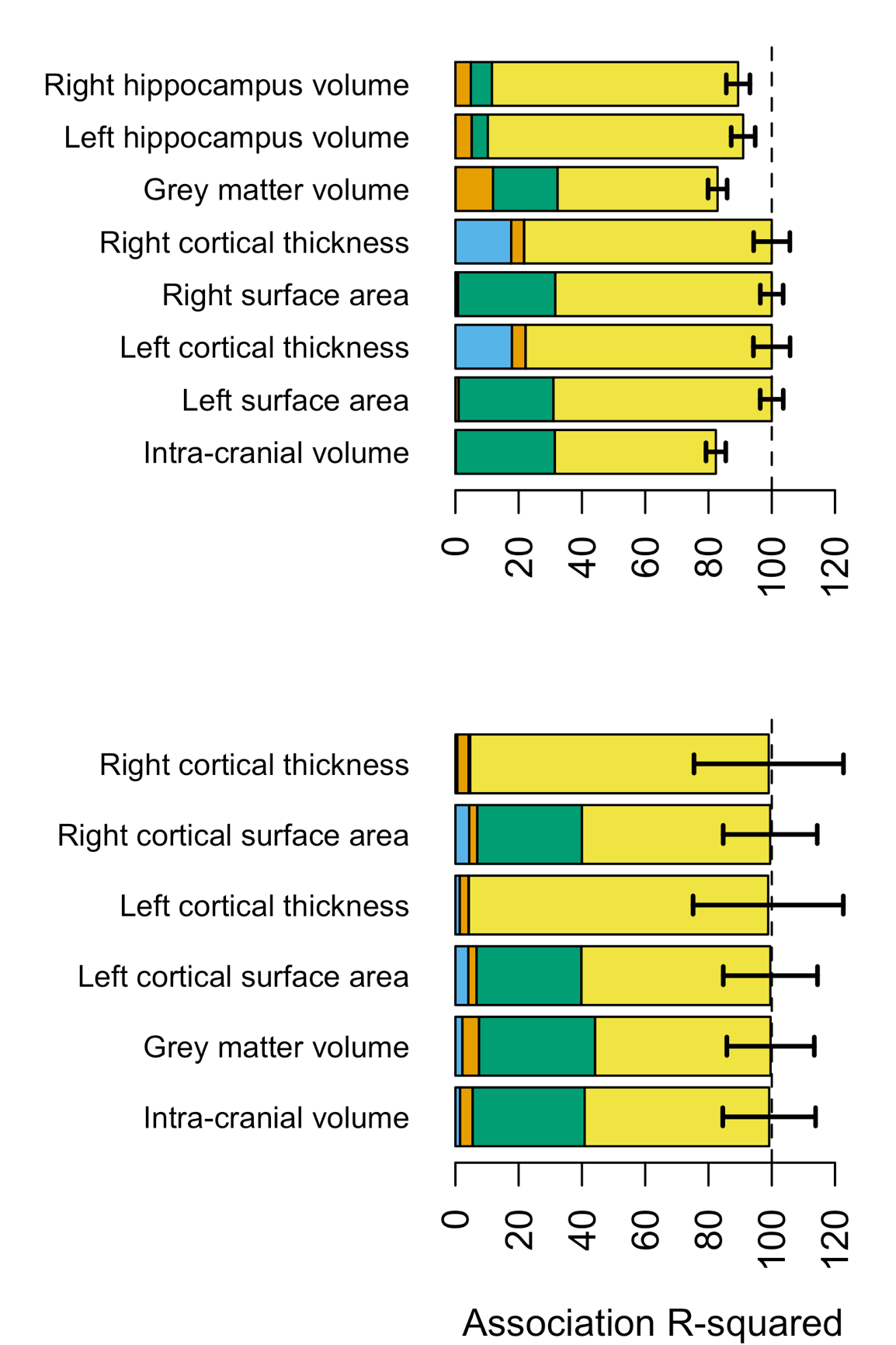
**Fig. S2. Morphometricity of brain phenotypes – positive controls**
As a positive control, we estimated the association between all grey-matter vertices and global measures of brain size, controlling for acquisition age and sex. Results are shown for the UKB discovery sample (top panel) and the HCP sample (bottom). We observe large associations (R2 in the range of 0.51-0.81 for the UKB, R2 in 0.55-0.94 for the HCP) resulting in more than 83% of the phenotypic variance (>99% for the HCP) accounted for when adding the contribution of the covariates. Note that in the UKB, we found an association between left and right average thickness and processing option (T1w vs. T1w +T2 FLAIR) – light blue bar: R2~0.20.

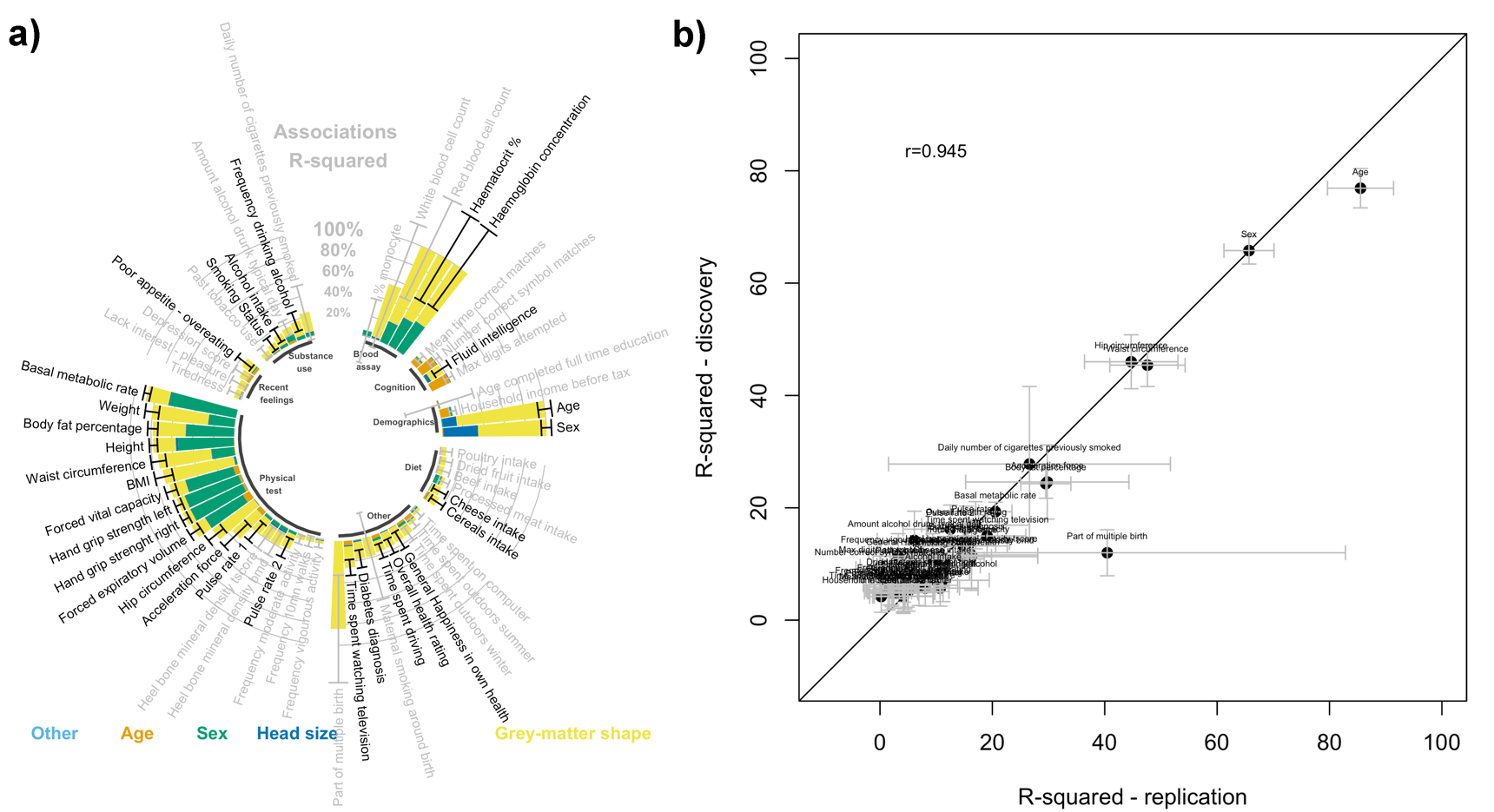

**Fig. S3: Morphometricity in the UKB replication sample under baseline covariates.**

Panel a) presents the summary of the replication analysis in the UKB. Only the 58 significant phenotypes from the UKB discovery sample were included in the analysis. All are shown here. Black bars correspond to 95% confidence intervals. Phenotypes showing significant morphometricity in the replication sample (p<0.05/58) are labelled in black. To note, the replication sample was only half the size of the discovery one, with a clear incidence on power. Blood assay variables and being part of multiple birth suffered from a small number of observations (N~300), which explains the large confidence intervals. Panel b) compares the replication morphometricity estimates (X axis) to the discovery ones (Y-axis). We excluded blood assay variables, for which the small N led to unstable parameter estimation. We found a great concordance of results between the 2 independent UKB samples as indicated by a correlation of 0.95 between the 2 sets of results.

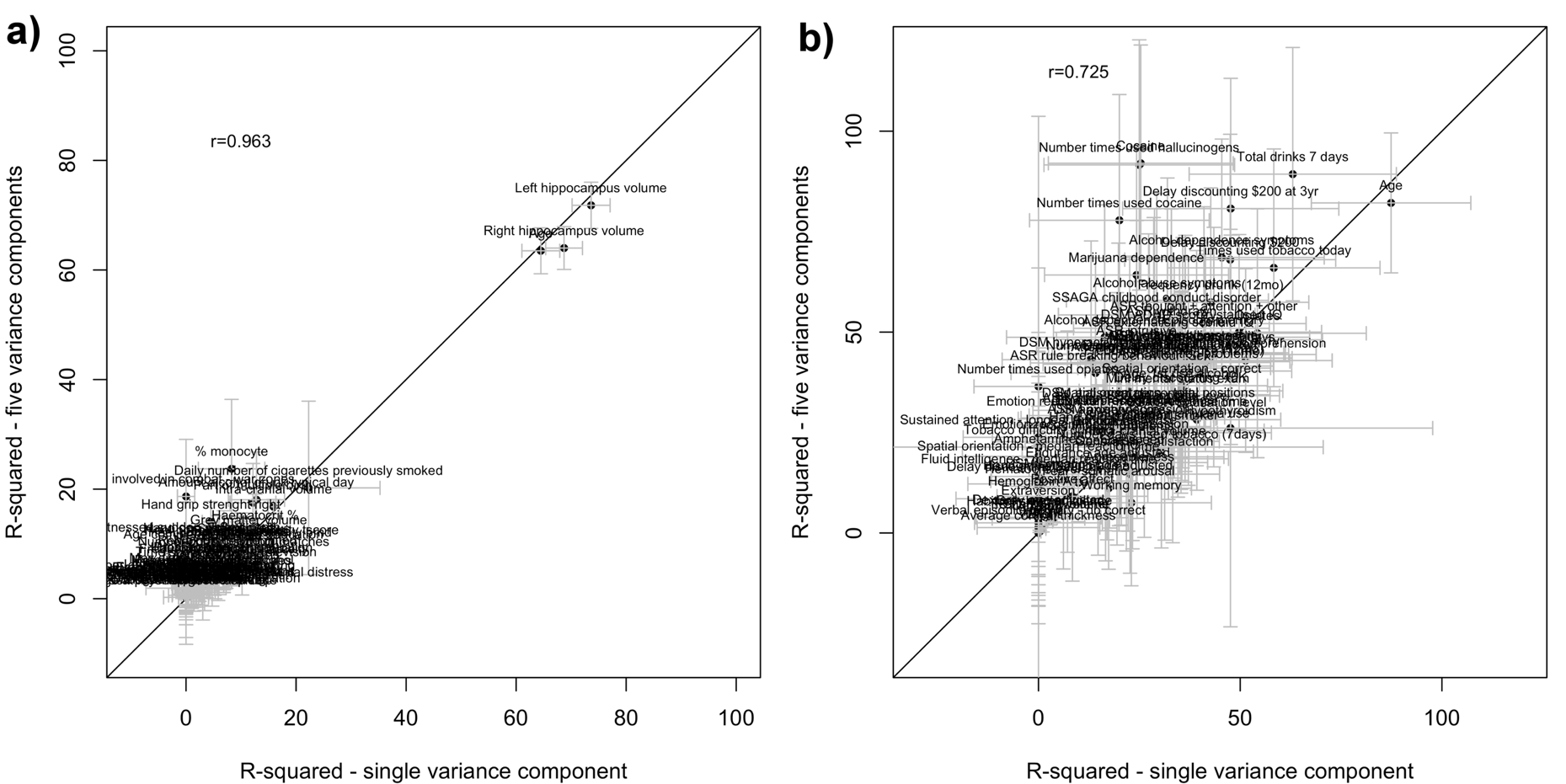

**Fig. S4: Scatterplots of association R^2^ from MLMs, comparing results when fitting a single BRM versus 4 BRMs corresponding to all different brain modalities (cortical thickness, cortical surface area, subcortical thickness and subcortical curvature).**

Panel (a) shows the results for the UKB sample, panel (b) shows the results for the HCP sample. Note that fitting multiple variance components comes at an increased computational cost (required memory increases with the number of BRM and each iteration of the REML algorithms often takes more time) and a slightly increased standard error of the estimate of the overall variance explained(12). In addition, when modelling 5 variance components, the AI-REML algorithm failed to converge for 58 UKB phenotypes and 71 HCP phenotypes because of the increased uncertainty in the estimate of variance explained by each component due to smaller number of brain measurements in an individual component in comparison with the total. The correlation between the association R^2^ from the two approaches appears on the plot.

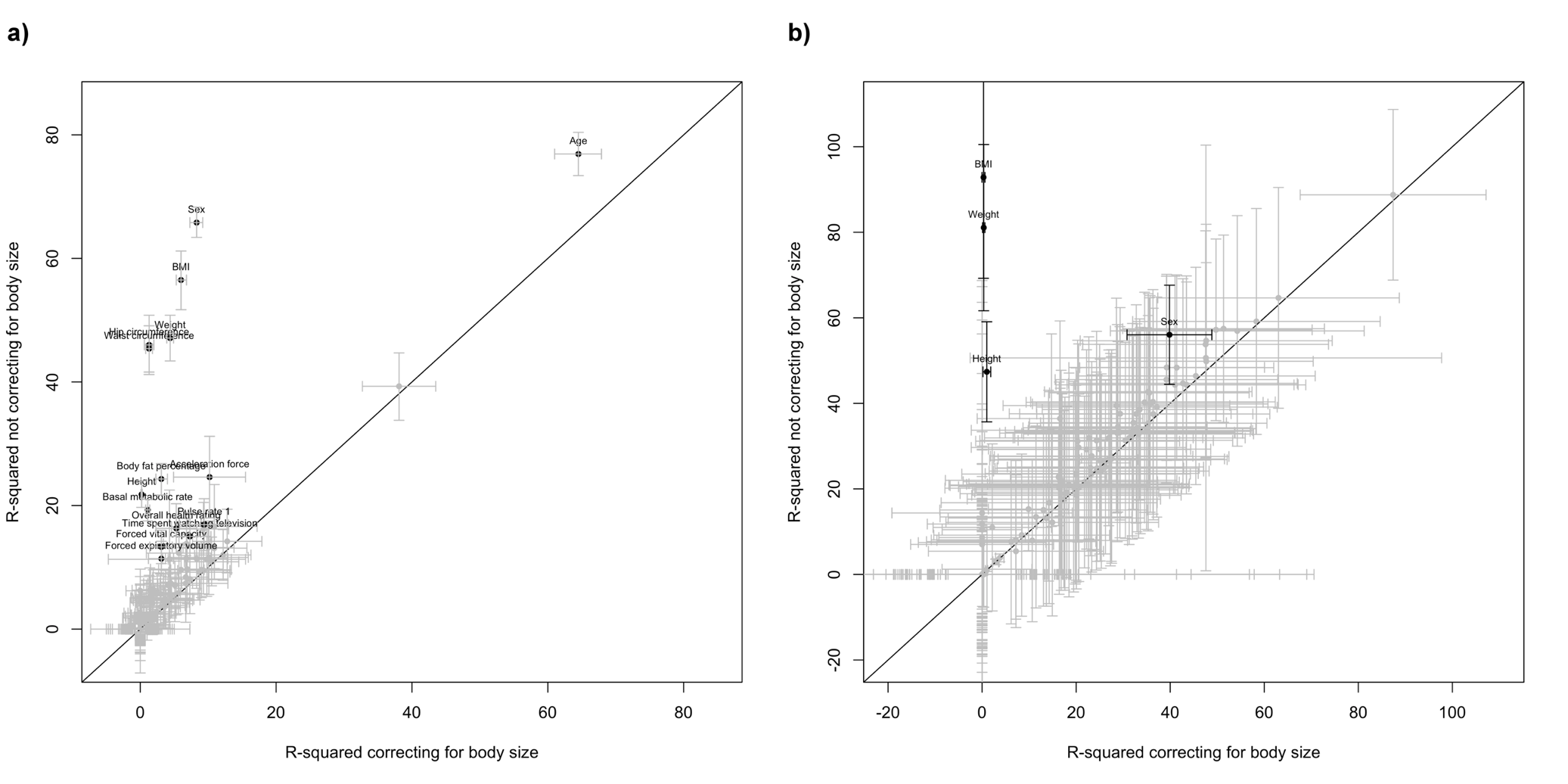

**Fig. S5: Effect of body size on phenome association with grey-matter shape. Scatterplots of association R^2^ for all the phenotypes before and after correcting for body-size variables.**
The bias and confound induced by body size can be visually appreciated, especially in the UKB discovery sample (panel (a)) and to a lesser extent in the HCP (panel (b)). Bars represent the 95% confidence intervals.

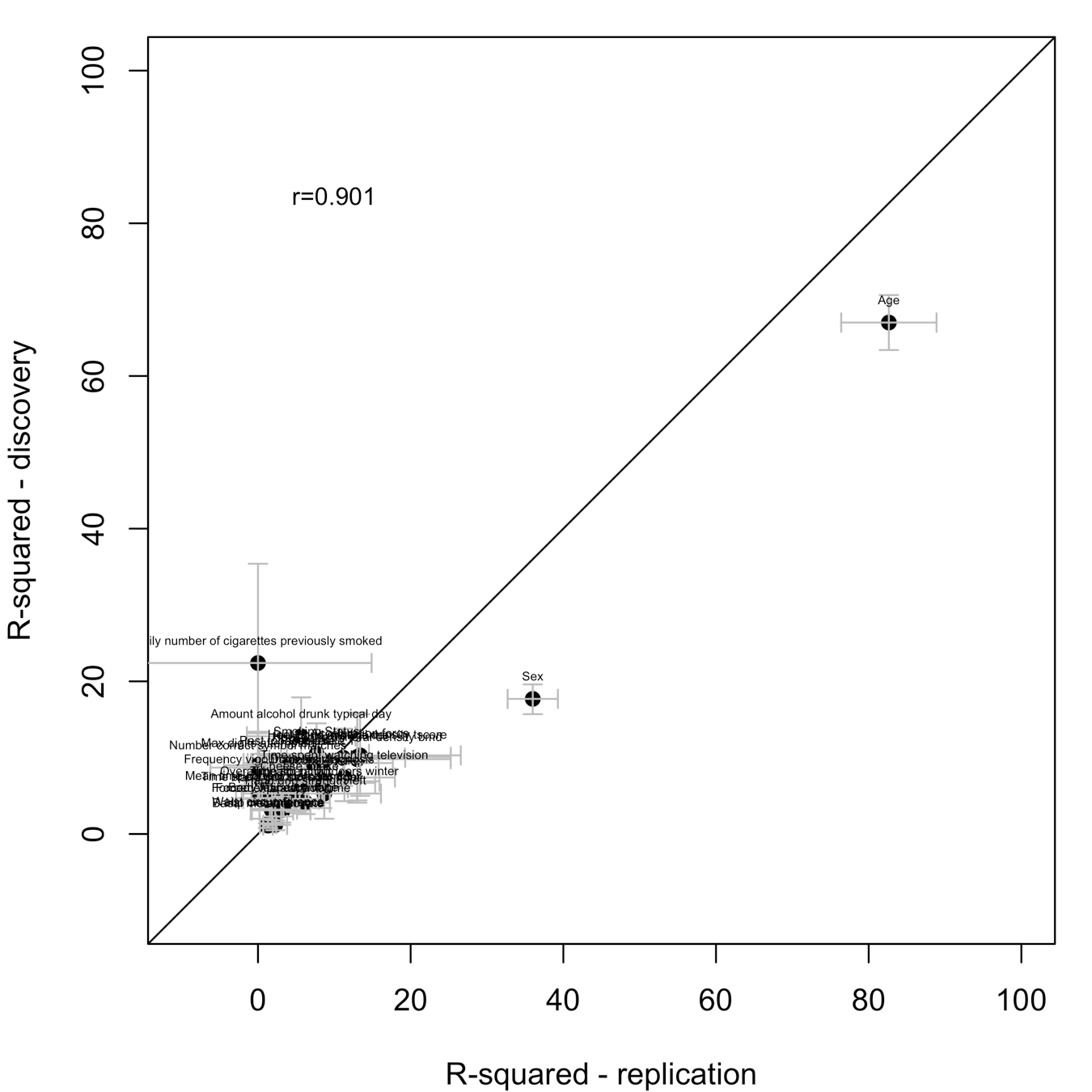

**Fig. S6: Morphometricity estimates correcting for height, weight and BMI in addition to baseline covariates, in the UKB replication sample (X-axis) and the UKB discovery sample (Y-axis).**
Only phenotypes displaying significant morphometricity in the UKB discovery set are included. Bars correspond to the 95% confidence intervals. Age and sex showed larger morphometricity in the replication sample. This was not due to different association strength with the covariates (see Fig. S1) and the demographic differences were limited (Dataset S1). We think that these could be true differences in morphometricity that might arise from outliers or heterogeneity in the discovery sample (composed of 3 imaging waves, with some marginal imaging protocol changes). In comparison, the replication sample, was a single wave of the UKB data collection.

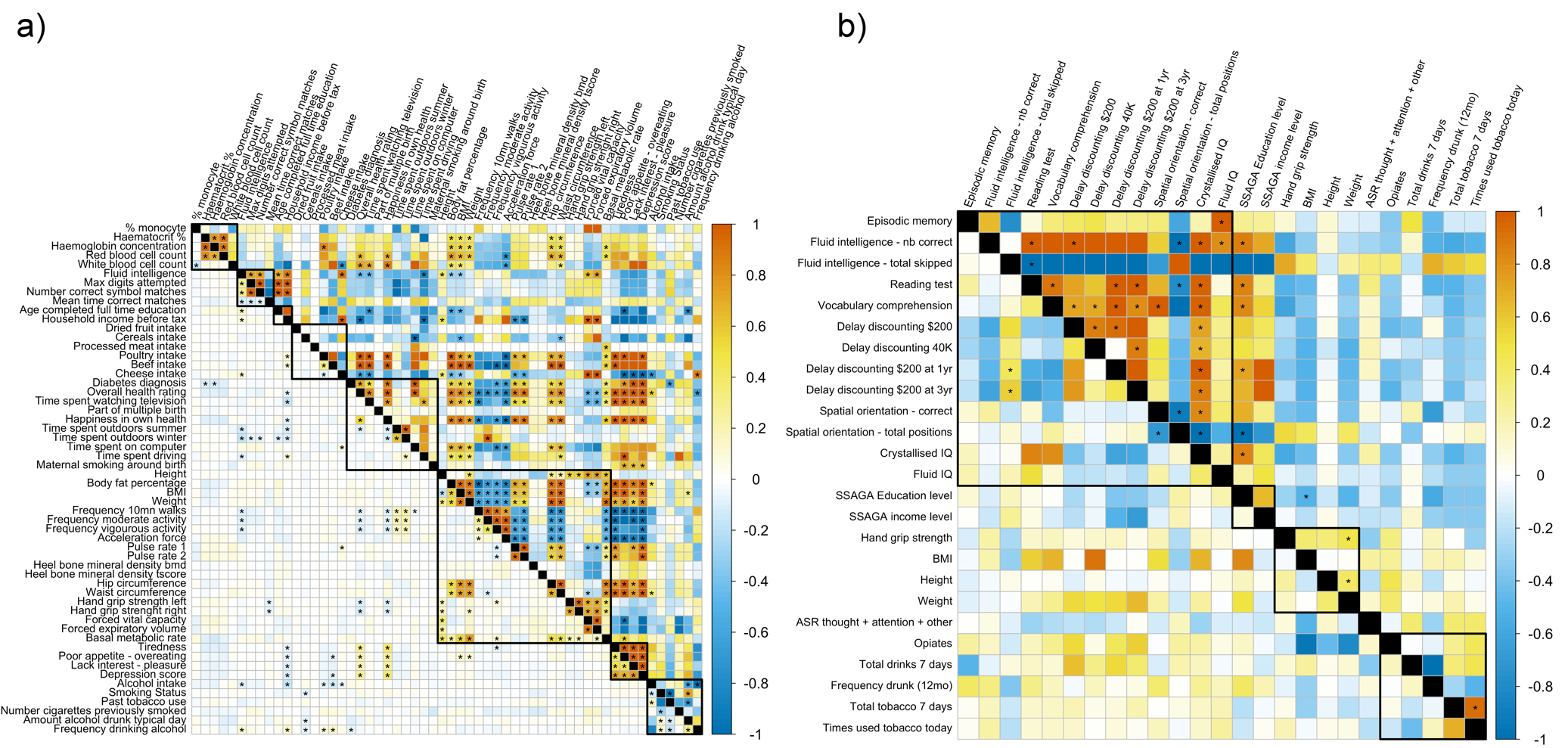

**Fig. S7: Grey-matter and residual correlations under the baseline model (i.e. not correcting for body size).**
Estimates are shown for the UKB (panel a) and HCP (b). Grey-matter correlation is shown above the diagonal, and residual correlation below the diagonal.

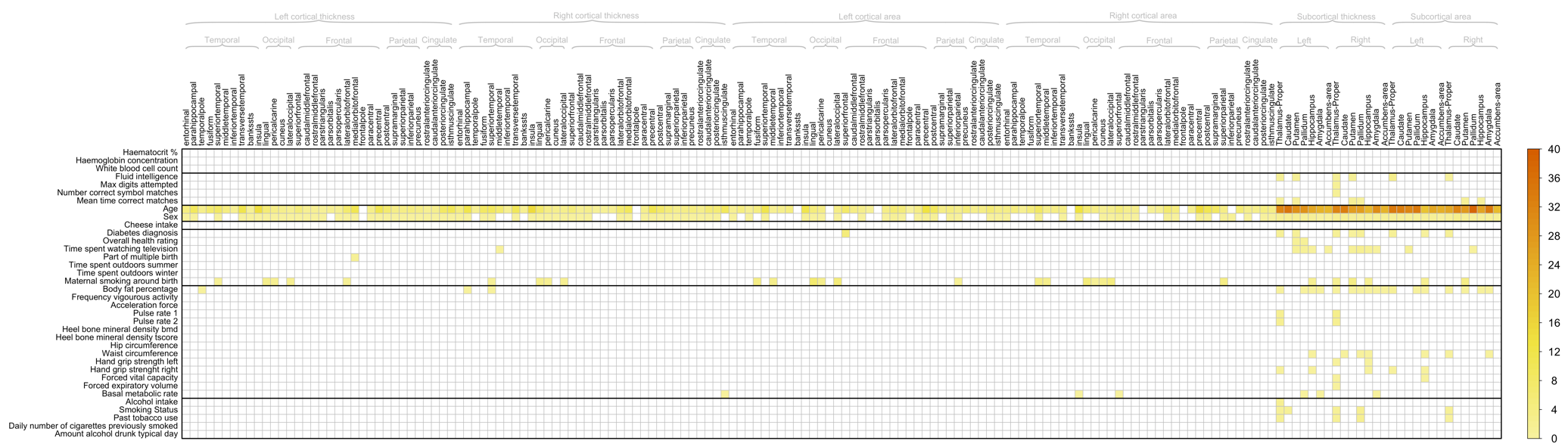

**Fig. S8: Region Of Interest (ROI) based LMMs in the UKB.**
Plot displays the significant association R^2^ between each UKB phenotype associated with grey-matter shape in Figure 1a and the grey-matter vertices from each of the Desikan (13) atlas ROI. Results include baseline covariates as well as height, weight and BMI.

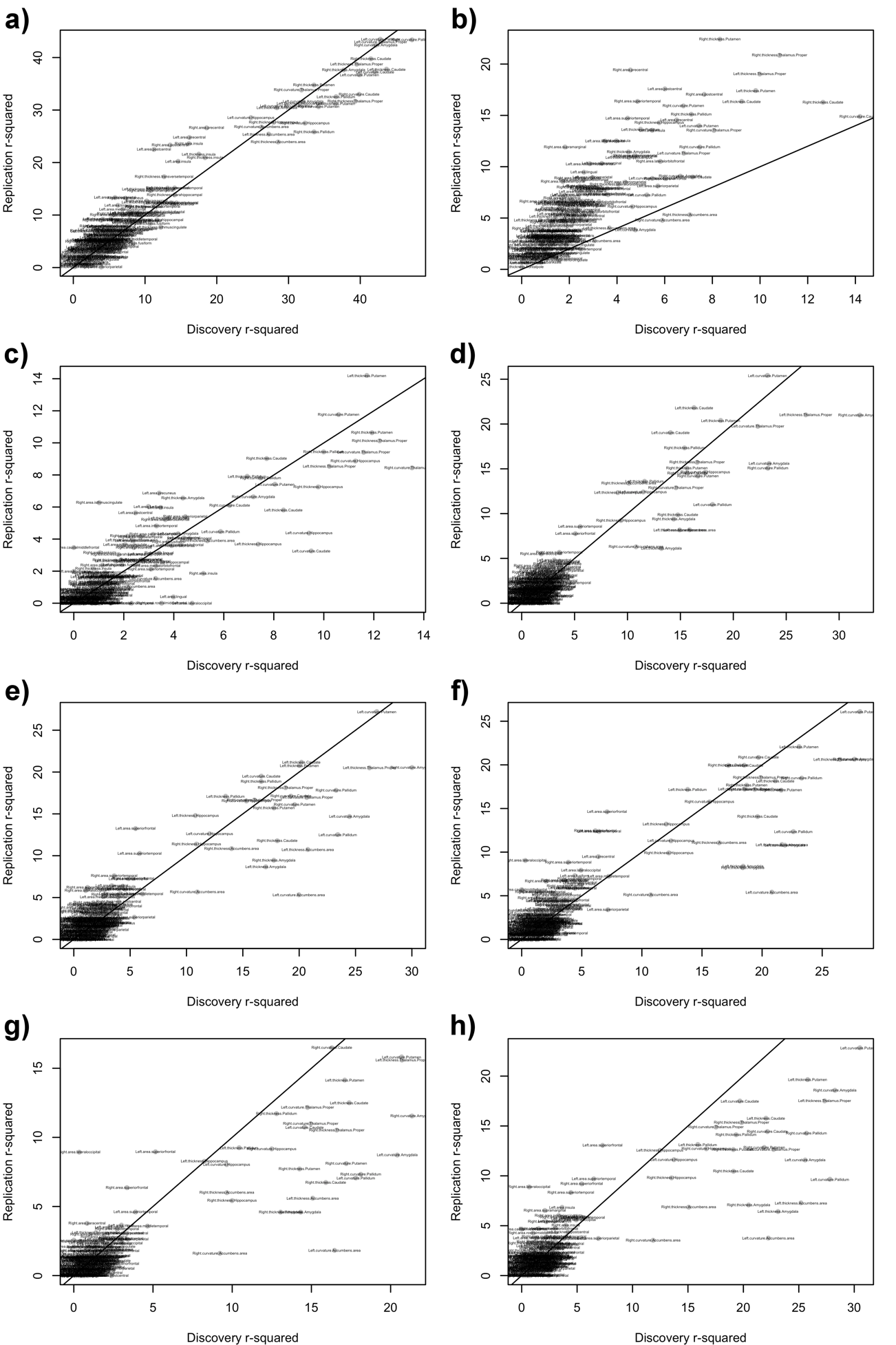

**Fig. S9: ROI based associations between UKB discovery and replication samples for selected phenotypes.**

Each panel correspond to a single phenotype, and displays the association R2 of this phenotype with each ROI of interest considered. The results found in the replication sample (Y-axis) are plotted as a function of the discovery results (X-axis). The labels indicate which ROI corresponds to which point in the figure. Panel a) Age; b) sex; c) height; d) body fat %; e) BMI; f) weight; g) hip circumference; h) waist circumference.

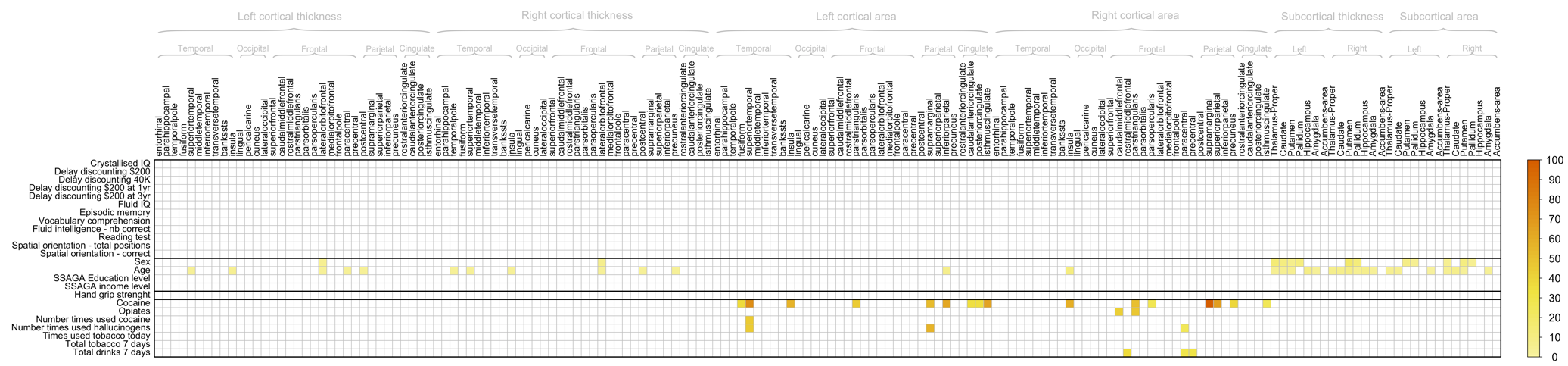

**Fig. S10: Region Of Interest (ROI) based MLMs in the HCP.**

Plot displays the significant association R^2^ between each UKB phenotype associated with grey-matter shape in Figure 1b and the grey-matter vertices from each of the Desikan atlas ROI.

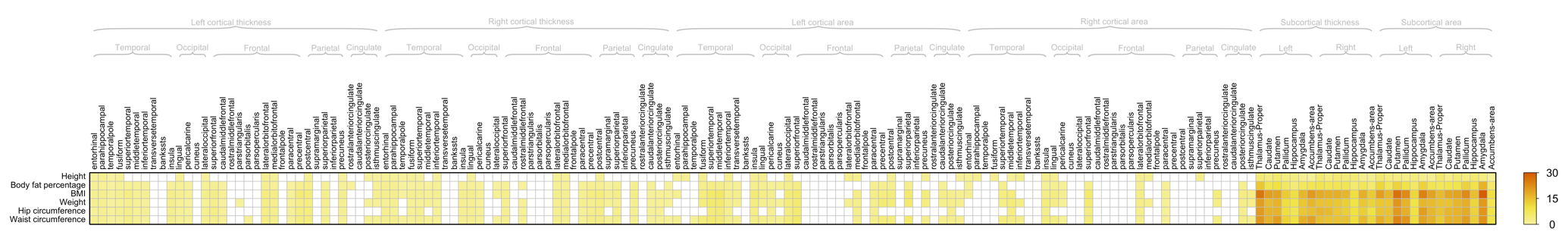

**Fig. S11: Region Of Interest (ROI) based LMMs in the UKB for body size variables**
Baseline covariates used.

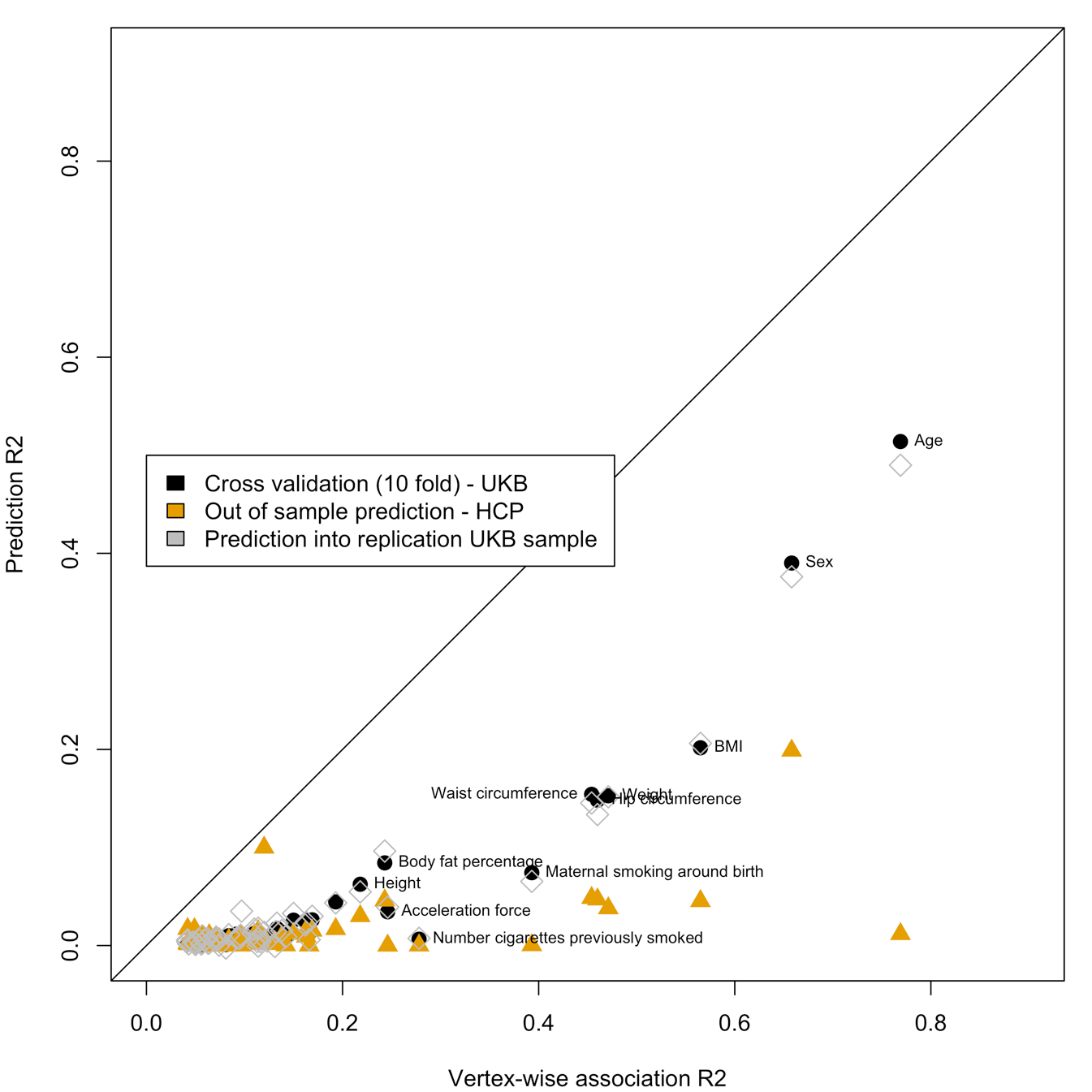

**Figure S12: In sample and out of sample prediction accuracy as a function of the total association R^2^ (baseline covariates)**
Labels highlight some of the significant prediction. As predicted by the theory, the prediction accuracy is capped by the total association R^2^ (points below the diagonal). In addition, out of sample prediction results in a lower prediction accuracy than in-sample prediction.

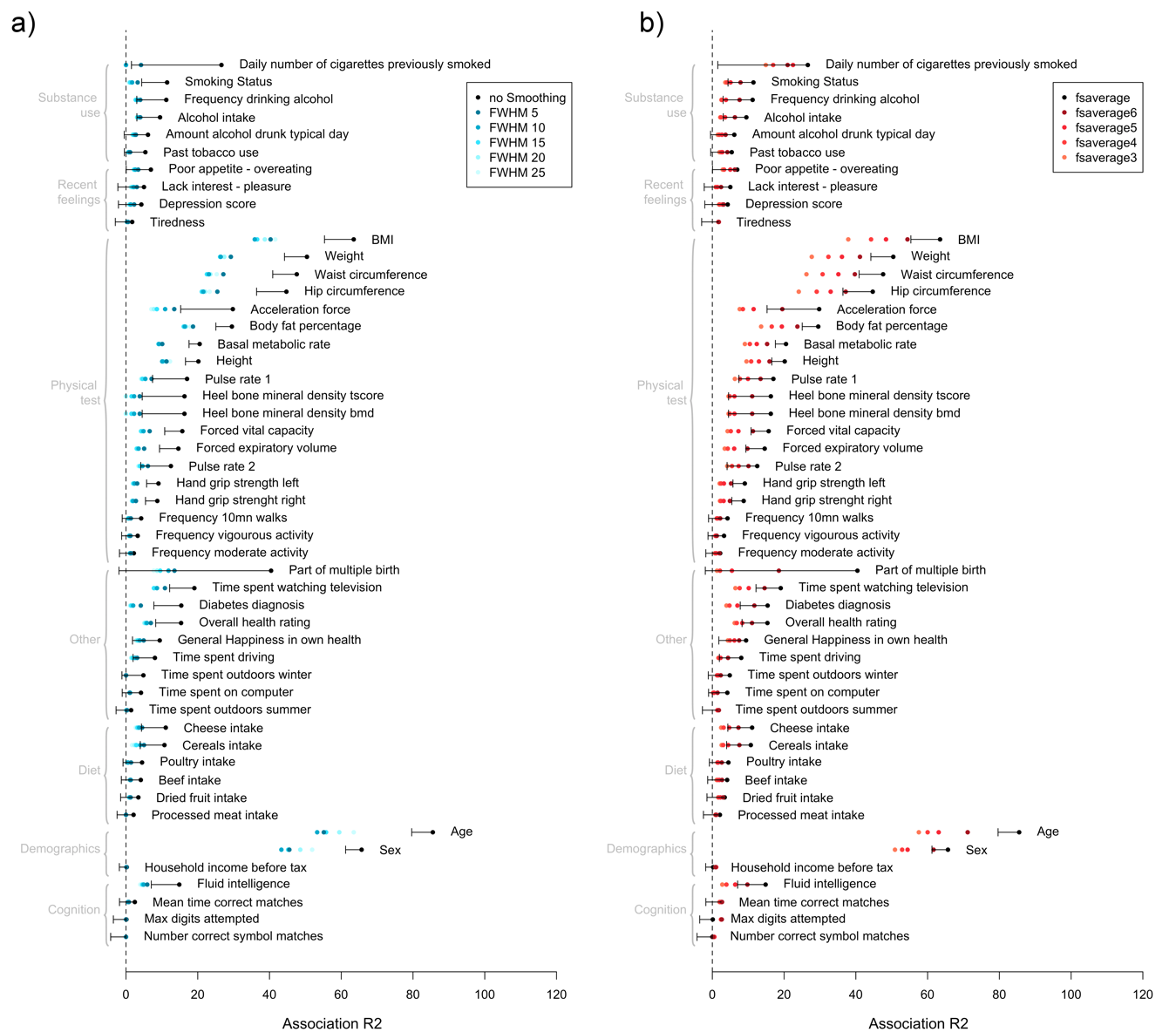

**Figure S13: Effect of cortical mesh smoothing and mesh choice on the brain-morphometricity estimates (UKB replication samples.**

Blood assay Variables (with n<500 observations) were excluded from the analysis due to unstable estimates and large SE.

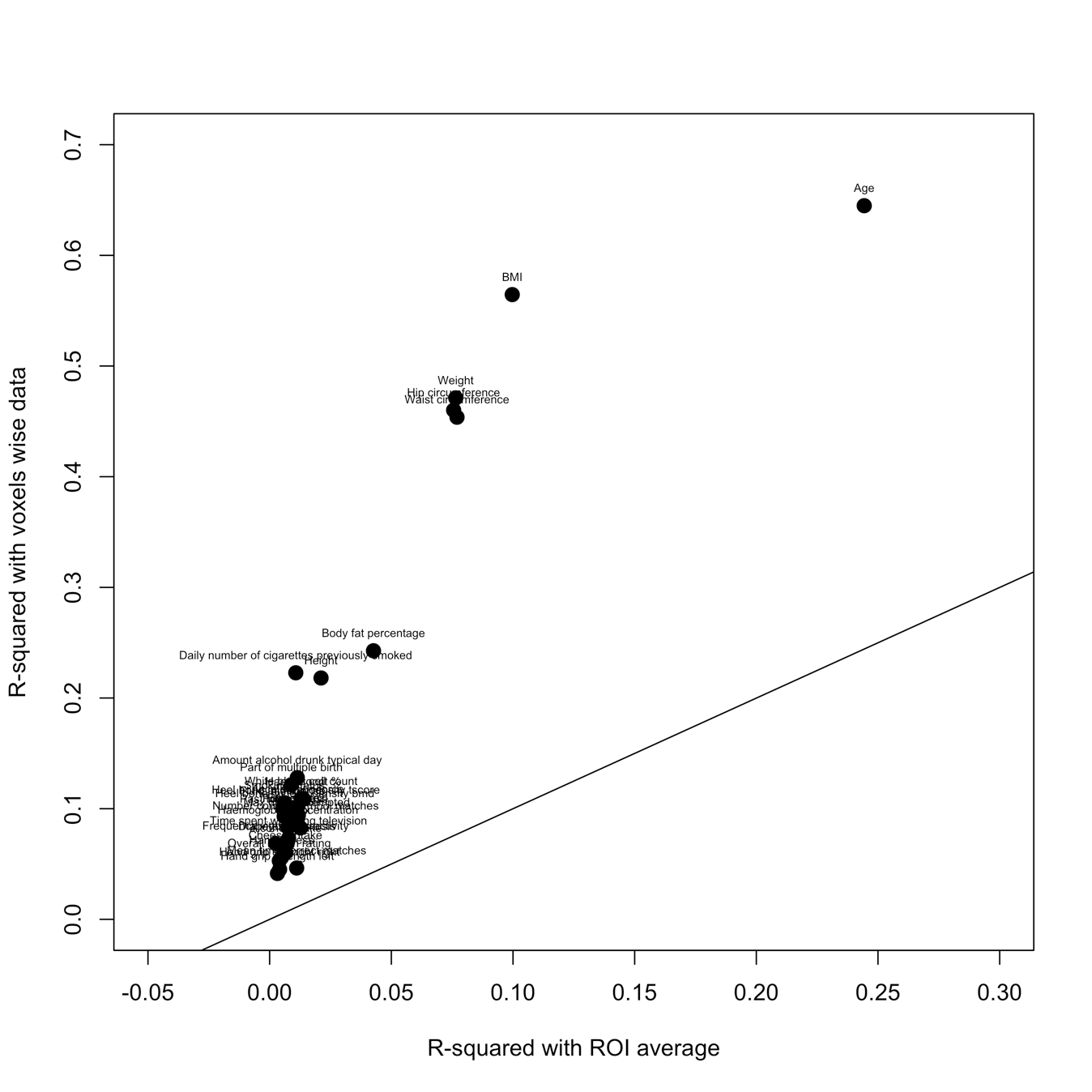

Fig. S14: Scatter plot comparing, for each UKB phenotype, our association R^2^ from vertex-wise processing and that obtained from standard ENIGMA ROI based processing
See (<http://enigma.ini.usc.edu/protocols/imaging-protocols/)>. All the points are above the diagonal indicating a greater amount of information retained using the vertex-wise processing compared to the ROI based dimension reduction approach. For example: R^2^_age_Vertex_=0.64 vs. R^2^_age_ROI_=0.24; R^2^_CigarettesPreviouslySmoked_Vertex_=0.22, R^2^_CigarettesPreviouslySmoked_ROI_=0.011, R^2^_AmountDrunkTypicalDay_Vertex_=0.13, R^2^_AmountDrunkTypicalDay_ROI_=0.011, and also R^2^_BMI_Vertex_=0.56 vs. R^2^_BMI_ROI_=0.10 (baseline covariates). This demonstrates the superiority of full resolution (vertex-wise) analyses of structural brain images, compared with an atlas based dimension reduction.

Table S1. Replication of grey-matter correlations identified in the UKB discovery sample

| **Variable 1** | **Variable 2** | **Discovery** | **Replication** | | |
| --- | --- | --- | --- | --- | --- |
|  |  | **rGM** | **rGM** | **SE** | **pvalue** |
| Fluid intelligence | Max digits attempted | 0.71 | 1.0 | 7.3 | 0.29469 |
| Fluid intelligence | Number correct symbol matches | 0.72 | 1.0 | 25.1 | 0.31246 |
| Max digits attempted | Number correct symbol matches | 1.0 | 1.0 | 25.1 | 0.31246 |
| Fluid intelligence | Cheese intake | 1.0 | 0.82 | 0.2 | 0.0015689 |
| Time spent outdoors summer | Time spent outdoors winter | 1.0 | 1.0 | 0.2 | 0.1928 |
| Time spent watching television | Body fat percentage | 0.73 | 0.81 | 0.2 | 4.42E-05 |
| Frequency vigorous activity | Acceleration force | 1.0 | 0.62 | 0.4 | 0.13604 |
| Overall health rating | Pulse rate 1 | 1.0 | 1.0 | 0.5 | 0.01224 |
| Overall health rating | Pulse rate 2 | 1.0 | 1.0 | 0.5 | 0.012028 |
| Pulse rate 1 | Pulse rate 2 | 0.99 | 1.0 | 0.01 | 0.00049312 |
| Overall health rating | Waist circumference | 1.0 | 0.39 | 0.4 | 0.19506 |
| Body fat percentage | Waist circumference | 0.52 | 0.45 | 0.2 | 0.02677 |
| Pulse rate 1 | Waist circumference | 0.67 | 0.31 | 0.3 | 0.1675 |
| Pulse rate 2 | Waist circumference | 0.73 | 0.56 | 0.3 | 0.047662 |
| Hand grip strength left | Hand grip strength right | 0.92 | 0.84 | 0.06 | 1.76E-07 |
| Body fat percentage | Forced vital capacity | -0.66 | -0.45 | 0.2 | 0.026155 |
| Hand grip strength right | Forced vital capacity | 0.69 | 0.70 | 0.2 | 0.0015424 |
| Fluid intelligence | Forced expiratory volume | 1.0 | 0.48 | 0.3 | 0.062686 |
| Hand grip strength right | Forced expiratory volume | 1.0 | 0.79 | 0.2 | 0.00044463 |
| Body fat percentage | Basal metabolic rate | -0.69 | -0.75 | 0.1 | 3.89E-06 |
| Pulse rate 1 | Basal metabolic rate | -0.55 | -0.46 | 0.2 | 0.038444 |
| Waist circumference | Basal metabolic rate | -0.75 | -0.55 | 0.2 | 0.0055741 |
| Smoking Status | Past tobacco use | -0.98 | -0.93 | 0.03 | 0.0011549 |
| Alcohol intake | Amount alcohol drunk typical day | -0.89 | -1.0 | 0.1 | 0.00042018 |
| Smoking Status | Amount alcohol drunk typical day | 0.71 | 1.0 | 0.2 | 0.0044935 |
| Past tobacco use | Amount alcohol drunk typical day | -0.64 | -1.0 | 0.8 | 0.091905 |

**Table S3:** Summary of the prediction accuracy (R^2^) of the BLUP grey-matter scores. We constructed BLUP scores for the 39 UKB variables showing significant morphometricity and evaluated their predictive power in the UKB (10 fold-cross validation) and HCP sample. When the phenotype corresponding to the grey-matter score was not available in the HCP, we chose the closest available (e.g. waist circumference grey-matter score evaluated against BMI). We evaluate the prediction accuracy by fitting GLM controlling for height, weight and BMI as well as for the baseline covariates (acquisition, age, sex and head size); except for (#) denoting associations not controlling for height, weight and BMI. Rows in bold indicate significant association after correcting for multiple testing (p<0.05/39=1.3e-3) both in and out of sample.

|  | **In sample prediction (UKB)** | | | | **Prediction into UKB replication** | | | | **Out of sample prediction (HCP)** | | | | |
| --- | --- | --- | --- | --- | --- | --- | --- | --- | --- | --- | --- | --- | --- |
|  | **r** | **pvalue** | **R^2^** | **AUC (SE)** | **r** | **pvalue** | **R^2^** | **AUC (SE)** | **HCP variable predicted** | **r** | **pvalue** | **R^2^** | **AUC (SE)** |
| Haemoglobin concentration | 0.05 | 1.6e-05 | 0.0025 |  | 0.035 | 5.5e-01 | 0.0013 |  | Hemoglobin A1C | 0.015 | 6.8e-01 | <0.001 |  |
| Haematocrit % | 0.077 | 1.4e-10 | 0.006 |  | 0.037 | 5.7e-01 | 0.0013 |  | Hematocrit level 1 | 0.061 | 2.1e-02 | 0.0037 |  |
| White blood cell count | 0.042 | 2.2e-03 | 0.0018 |  | 0.016 | 3.4e-01 | <0.001 |  | Hematocrit level 1 | -0.00051 | 9.8e-01 | <0.001 |  |
| Mean time correct matches | 0.046 | 2.0e-06 | 0.0021 |  | 0.053 | 2.9e-04 | 0.0028 |  | Crystallised IQ | -0.046 | 9.7e-02 | 0.0021 |  |
| Number correct symbol matches | 0.09 | 1.1e-13 | 0.0081 |  | 0.069 | 2.5e-04 | 0.0048 |  | Crystallised IQ | 0.064 | 2.2e-02 | 0.0041 |  |
| Max digits attempted | 0.091 | 2.9e-14 | 0.0083 |  | 0.083 | 1.1e-05 | 0.0069 |  | Crystallised IQ | 0.069 | 1.3e-02 | 0.0048 |  |
| Fluid intelligence | 0.077 | 4.1e-14 | 0.0059 |  | 0.11 | 7.2e-12 | 0.011 |  | Fluid IQ | 0.027 | 3.5e-01 | <0.001 |  |
| Age | 0.64 | 0.0e+00 | 0.41 |  | 0.68 | 0.0e+00 | 0.46 |  | Age | 0.15 | 3.1e-08 | 0.024 |  |
| Heel bone mineral density bmd | 0.065 | 2.1e-06 | 0.0042 |  | 0.058 | 2.8e-03 | 0.0034 |  | Age | -0.015 | 5.8e-01 | <0.001 |  |
| Heel bone mineral density tscore | 0.065 | 1.8e-06 | 0.0043 |  | 0.056 | 3.9e-03 | 0.0031 |  | Age | -0.015 | 5.9e-01 | <0.001 |  |
| Sex | 0.26 | 0.0e+00 | 0.067 | 0.58 (0.0059) | 0.33 | 9.8e-305 | 0.11 | 0.8 (0.0064) | Sex | -0.25 | 8.0e-42 | 0.061 | 0.68 (0.016) |
| Cheese intake | 0.061 | 2.0e-09 | 0.0037 |  | 0.076 | 1.9e-07 | 0.0058 |  | SSAGA Education level | 0.029 | 3.2e-01 | <0.001 |  |
| Part of multiple birth | 0.078 | 4.1e-14 | 0.0061 | 0.66 (0.022) | 0.13 | 1.5e-03 | 0.016 | 0.72 (0.065) | Being a twin | 0.31 | 1.1e-28 | 0.098 | 0.69 (0.016) |
| Body fat percentage | 0.29 | 0.0e+00 | 0.085 |  | 0.31 | 7.7e-190 | 0.095 |  | BMI | 0.21 | 5.6e-13 | 0.045 |  |
| Waist circumference | 0.39 | 0.0e+00 | 0.16 |  | 0.38 | 2.0e-205 | 0.14 |  | BMI | 0.21 | 3.5e-13 | 0.046 |  |
| Frequency vigourous activity | 0.044 | 9.4e-06 | 0.002 |  | 0.033 | 1.6e-02 | 0.0011 |  | BMI | -0.0015 | 6.7e-01 | <0.001 |  |
| BMI | 0.45 | 0.0e+00 | 0.2 |  | 0.45 | 7.4e-235 | 0.20 |  | BMI | 0.21 | 2.4e-12 | 0.042 |  |
| Diabetes diagnosis | 0.062 | 4.7e-10 | 0.0038 | 0.6 (0.014) | 0.085 | 2.5e-09 | 0.0073 | 0.63 (0.019) | BMI | 0.0055 | 1.0e-01 | <0.001 |  |
| Overall health rating | 0.052 | 8.2e-08 | 0.0027 |  | 0.062 | 7.8e-06 | 0.0039 |  | BMI | -0.002 | 5.5e-01 | <0.001 |  |
| Time spent watching television | 0.083 | 3.0e-17 | 0.0068 |  | 0.12 | 1.8e-17 | 0.014 |  | BMI | -0.0025 | 4.5e-01 | <0.001 |  |
| Basal metabolic rate | 0.029 | 1.0e-24 | 0.00082 |  | 0.031 | 3.7e-14 | <0.001 |  | BMI | 0.0046 | 1.7e-01 | <0.001 |  |
| Hip circumference | 0.38 | 0.0e+00 | 0.15 |  | 0.36 | 7.3e-143 | 0.13 |  | BMI | 0.21 | 5.2e-13 | 0.045 |  |
| Time spent outdoors summer | 0.047 | 2.1e-06 | 0.0022 |  | 0.033 | 1.3e-02 | 0.0011 |  | BMI | -0.006 | 7.4e-02 | <0.001 |  |
| Time spent outdoors winter | 0.028 | 3.8e-03 | 0.00077 |  | 0.036 | 7.9e-03 | 0.0013 |  | BMI | -0.0073 | 3.0e-02 | <0.001 |  |
| Pulse rate 1 | 0.1 | 2.5e-23 | 0.01 |  | 0.13 | 4.0e-13 | 0.017 |  | BMI | 0.00012 | 9.7e-01 | <0.001 |  |
| Pulse rate 2 | 0.091 | 2.5e-19 | 0.0083 |  | 0.1 | 9.0e-09 | 0.0099 |  | BMI | -0.00057 | 8.7e-01 | <0.001 |  |
| Height | 0.25 | 6.5e-318 | 0.062 |  | 0.23 | 2.6e-132 | 0.054 |  | Height | 0.17 | 1.8e-17 | 0.03 |  |
| Acceleration force | 0.07 | 7.5e-08 | 0.0049 |  | 0.098 | 4.1e-06 | 0.0096 |  | Hand grip strength | 0.014 | 4.5e-01 | <0.001 |  |
| Hand grip strenght right | 0.039 | 1.8e-09 | 0.0015 |  | 0.052 | 1.4e-08 | 0.0027 |  | Hand grip strength | -0.0051 | 7.9e-01 | <0.001 |  |
| Forced vital capacity | 0.03 | 7.4e-06 | 0.00092 |  | 0.045 | 1.1e-05 | 0.002 |  | Hand grip strength | 0.0081 | 6.7e-01 | <0.001 |  |
| Forced expiratory volume | 0.027 | 2.3e-04 | 0.00072 |  | 0.06 | 7.8e-08 | 0.0036 |  | Hand grip strength | 0.019 | 3.3e-01 | <0.001 |  |
| Hand grip strength left | 0.039 | 1.6e-09 | 0.0015 |  | 0.064 | 2.9e-12 | 0.0041 |  | Hand grip strength | -0.019 | 3.1e-01 | <0.001 |  |
| Weight | 0.39 | 0.0e+00 | 0.15 |  | 0.39 | 5.8e-231 | 0.15 |  | Weight | 0.19 | 1.2e-12 | 0.036 |  |
| Alcohol intake | 0.074 | 1.0e-13 | 0.0055 |  | 0.096 | 2.9e-11 | 0.0092 |  | Frequence alcohol use (12mo) | 0.06 | 4.3e-02 | 0.0036 |  |
| Amount alcohol drunk typical day | 0.063 | 5.9e-08 | 0.0039 |  | 0.075 | 6.1e-06 | 0.0056 |  | Frequence alcohol use (12mo) | -0.0055 | 8.5e-01 | <0.001 |  |
| Past tobacco use | 0.043 | 1.7e-05 | 0.0019 |  | 0.076 | 2.2e-07 | 0.0057 |  | FTND score | 0.024 | 6.8e-01 | <0.001 |  |
| Smoking Status | 0.06 | 2.6e-09 | 0.0036 |  | 0.1 | 2.5e-12 | 0.01 |  | FTND score | -0.013 | 8.3e-01 | <0.001 |  |
| Number cigarettes previously smoked | 0.066 | 1.0e-03 | 0.0043 |  | 0.053 | 4.9e-02 | 0.0028 |  | FTND score | 0.011 | 8.5e-01 | <0.001 |  |
| Maternal smoking around birth | 0.26 | 9.8e-132 | 0.069 | 0.66 (0.0067) | 0.25 | 1.7e-08 | 0.063 | 0.65 (0.027) | FTND score | 0.19 | 8.9e-04 | 0.037 |  |

**Dataset S1:** Descriptive table of the UKB variables used in the analysis (for the discovery and replication UKB sets). Comparison between final sample participants excluded from the analysis due to failed processing and QC. UKB discovery and replication samples used in the analyses are further compared.

**Dataset S2:** Descriptive table of the HCP variables used in the analysis and comparison with participants excluded from the analysis due to QC.

**Dataset S3:** Detailed results of variance components analysis in the UKB (discovery and replication samples). Includes results presented in **Figure 1a**. Results include fixed effect associations (associations with covariates) as well as morphometricity. Baseline covariates are used in the first part of the table but results after accounting for height, weight and BMI are also presented.

**Dataset S4:** Detailed results of variance components analysis in the HCP. Results of **Figure 1b.**

**Dataset S5:** Grey-matter and residual correlations in the UKB. Table of the values presented in **Figure 2a** (UKB). The table contains correlations and p-values of the phenotypic and brain correlations. Format: r; (SE; p-value).

**Dataset S6:** Grey-matter and residual correlations in the HCP. Table of the values presented in **Figure 2b** (HCP). The table contains correlations and p-values of the phenotypic and brain correlations. Format: r; (SE; p-value).

**Dataset S7:** Association R^2^ between phenotypes and each ROI in the UKB discovery sample. Data is shown under the form of %variance (SE), p-value. This data corresponds to the **Figure S8**. Covariates include baseline + body size (height, weight and BMI).

**Dataset S8:** Association R^2^ between body size phenotypes and each ROI in the UKB discovery sample. Data is shown under the form of %variance (SE), p-value. This data corresponds to the **Figure S10**. Baseline covariates only.

**Dataset S9:** Association R^2^ between phenotypes and each ROI in the UKB replication sample. Data is shown under the form of %variance (SE), p-value.

**Dataset S10:** Association R^2^ between phenotypes and each ROI in the HCP sample. Data is shown under the form of %variance (SE), p-value. This data corresponds to the Supplementary **Figure S11**.

**Dataset S11:** Accuracy from BLUP scores achieved using baseline covariates only. Results include 10-fold cross validation (UKB discovery), as well as prediction into UKB replication and HCP. These results are summarized in **Figure S11**.

**Dataset S12:** Brain-morphometricity results varying coarseness of cortical meshes in the UKB discovery sample. Significant results (from **Figure 1)** are presented in **Figure 4**.

**Dataset S13:** Brain-morphometricity results varying smoothing and coarseness of cortical meshes in the UKB replication sample. All UKB phenotypes are included. Significant results (from **Figure 1)** are presented in **Figure S13**.
